# User guide to the red blood cell model (RCM), a multiplatform JAVA-based model of human red blood cell homeostasis

**DOI:** 10.1101/2020.03.07.981779

**Authors:** Simon Rogers, Virgilio L. Lew

## Abstract

We introduce here a new multiplatform JAVA-based mathematical-computational model of RBC homeostasis for investigating the dynamics of changes in RBC homeostasis in health and disease. We provide a brief overview on the homeostasis of human RBCs and on the general biophysical principles guiding the modelling design. By way of a detailed tutorial we apply the model to investigate in depth the multiple effects associated with RBC dehydration induced by potassium permeabilization, a necessary preliminary for understanding the pathophysiology of a wide group of inherited haemolytic anaemias, a subject of intense current research and clinical interest. Using the red cell model (RCM), we design and run in silico representations of experimental protocols to study global RBC responses to calcium and potassium permeabilization covering a wide range of experimental, physiological and pathological conditions. Model outputs report the evolution in time of all the homeostatic variables in the system allowing, for the first time, a detailed and comprehensive account of the complex processes shaping global cell responses. Analysis of the results explains the mechanisms by which the entangled operation of all the RBC components link cell dehydration and protein crowding to cell acidification and to the generation of hypertonic, alkaline effluents. Open access to the RCM in a GitHub repository, together with the tutorial primed for a specific investigation pave the way for researchers and clinicians to apply the model on many different aspects of RBC physiology and pathology.

## The red blood cell model (RCM) and User Guide: what to expect

The User Guide contains example-based tutorials on how to apply the RCM in research, clinical and teaching contexts. Parameter values in the model are tightly constrained by solid data from the literature enabling model predictions to be semi-quantitative to levels of between 5 and 20%. To interpret this margin it is important to realize a critical difference between experiments at the bench and their model representation. At the bench, experiments are performed on cell populations in suspensions or on chosen isolated cells from a cell population sample, as with patch-clamp or electrode measurements. The RCM, on the other hand, predicts the behaviour of a virtual cell clone defined by the constitutive properties attributed to the modelled cell when specifying initial conditions.

The model becomes a singularly useful tool when comparisons between predicted and measured results depart from that accuracy margin. Such departures expose either ignored model components required for a particular cell response, or unexpected abnormalities or heterogeneities in the RBC population under investigation. To investigate population behaviour with the model it is necessary to run multiple identical simulations on clones defined with hypothesized variations in cell properties or components [1, 2]. Weighted proportions attributed to each variant within a defined distribution render predicted outcomes for the integrated response of the cell population under study. These can then be compared with experimental results to trace the nature of the hypothesized variant or distribution abnormality. The full sets of red blood cell parameters and variables are listed in the Appendix.

## General Introduction

### A brief primer on red blood cell homeostasis

The main function of RBCs is to mediate the transport of O_2_ and CO_2_ between lungs and tissues by mechanisms evolved to minimize the energy cost to the organism. A first critical feature enabling such economy is the extremely low cation permeability of the RBC membrane [3–5]. This allows the cells to maintain steady volumes for extended periods of time with minimal cation traffic, pump-leak turnover rates and ATP consumption. Glycolytic ATP turnover by the full RBC mass of a healthy human adult amounts to less than 0.06% of the total body ATP turnover [6].

A second critical feature of the optimized economy concerns the compromise between RBC turnover rate and circulatory lifespan. RBCs are the most abundant cells in the body, their mass adapted for adequate gas transport at all levels of physiological demand. Biosynthetic and biodegradable replacement of such a large cell mass imposes a heavy metabolic cost to the organism which can only be reduced by extending the circulatory lifespan of the cells thereby reducing their replacement frequency. Circulatory longevity, on the other hand, is limited by the extent to which RBCs, without nucleus and organelles, and devoid of biosynthetic capacity, can sustain the functional competence of its metabolic and membrane transport components required for volume stability and optimal rheology. Optimal rheology requires that the RBCs retain a large degree of deformability for rapid passage through narrow capillaries and for ensuring minimal diffusional distances for gas-exchange across capillary walls. Deformability, in turn, depends on the RBCs maintaining their volume well below their maximal spherical volume (reduced volume), a condition fulfilled when their reduced volume is kept within a narrow margin around 60% [6–8]. For human RBCs with a mean circulatory lifespan of about 120 days, this represents a substantial challenge.

RBC homeostasis involves a subset of basic cellular processes that attempt to maintain and restore cell volume and integrity throughout all dynamic changes elicited by physiological stress. Although all RBC components participate in global cell responses, the main players within the subset concerned with RBC homeostasis are the full constellation of passive and active membrane transporters, haemoglobin as the main macromolecular colloidosmotic contributor, proton and calcium buffer, the variety of impermeant cell metabolites and solutes contributing fixed negative charges and additional proton, calcium and magnesium buffering capacity to the RBC cytoplasm. Most of the haemolytic anaemias affecting humans result from parasitic invasions or from inherited mutations in components of the homeostasis subset. Early versions of the red blood cell model proved a valuable tool for unravelling the complex pathophysiology in some of these diseases [9–12].

### Introducing the red blood cell model

The RCM describes the dynamic behaviour of a suspension of red blood cell clones in plasma-like media, treated as a closed two-compartment system. The physical laws constraining the behaviour of such a system are charge and mass conservation.

Models of cellular homeostasis start with an electroneutral cell system in an initial reference steady-state or quasi-steady-state. Following perturbations, the equation that ensures sustained charge conservation and electroneutrality in a system with i-ion components throughout all dynamic changes is ∑ Ii = 0. Each Ii describes the current carried by individual ions through channels, carriers and pumps across the membrane. The relation between currents and individual ion fluxes, Fi, is given by ∑Ii = *F*\*∑zi*Fi, where zi in the valence of ion i and *F* is the Faraday constant. ∑Ii is a function of the membrane potential, Em, of the concentration of the transported substrates on each membrane side, on modulating factors such temperature, pH, [Ca^2+^]_i_, [Mg^2+^]_i_, other ions, solutes and metabolites. ∑Ii = 0 is therefore a complex equation. When the parameters and kinetics of all the individual Fi equations are either known or phenomenologically defined, as expected for a system mature for modelling such as the human red blood cell, ∑Ii = 0 becomes an implicit equation in Em, the single unknown left.

We apply in house solution routines for the ordinary differential equations governing the dynamic behaviour of the system and also our own developed cord-approximation Newton-Raphson routine for the solution of all implicit equations in the model. Extensive application of these operators in different cell systems confirmed their superior performance over alternatives on code-economy and speedy convergence to solutions.

#### Avoidance of explicit equations for Em in homeostasis models

Models seeking simplified explicit formulations of the Em equation, as attempted in the past [13, 14], are misguided. To retain the flexibility to change the flux-kinetics of individual transporters in the search for improved fits between predicted and experimental results, the freedom to modify the terms within ∑ Ii = 0 is essential, a process fatally compromised by seeking simplifying assumptions in the search for explicit Em formulations.

#### Membrane capacitance may be neglected in homeostasis models

Em variations change the charge on the membrane capacitor. Such changes can almost always be ignored because the amount of charge displaced, as well as the duration of the current transients, Ic(t) = C*dV/dt, are orders of magnitude below the magnitude and time-course of the ionic currents of homeostatic relevance [15]

#### ∑ Ii = 0 and Entanglement

Charge conservation imposes an extraordinary level of entanglement in the processes that shape global cell responses. For example, consider an ion channel becoming activated in a cell at rest. Em changes and so do all voltage-sensitive Ii, causing secondary changes in the concentrations of other transported substrates. Some of these substrates are shared by electroneutral transporters, altering their fluxes and causing additional changes in the concentrations of intracellular solutes. In turn, all transport, biochemical and metabolic processes that depend on the concentration of intracellular solutes and on cell osmolarity also become affected generating chains of complex, interlinked interactions all influencing the global cell response. The magnitude of individual and global changes may vary between miniscule or huge scales, but the web of interconnected influences is always there. Homeostatic entanglement renders intuition a very fallible instrument for predicting global cell behaviour and for understanding the mechanisms behind complex cell responses, as was amply demonstrated in the past [10, 11, 16–24].

#### The JAVA version of the RCM

Building on the same approach in which unexpected but experimentally verified model predictions helped solve issues of substantial relevance in biology and medicine in the past [9, 25–31] we introduce here a vastly extended and fully updated JAVA model of the homeostasis of human RBCs using the Gardos effect, of historic and renewed relevance, to demonstrate how to use the model to unravel the molecular interactions that participate in processes controlling or affecting RBC hydration in physiological and pathological conditions and to explain the mechanisms behind, an issue of intense current haematological interest [8, 32–39] .

The JAVA model introduced here includes a graphical user interface along with a link to the GitHub repository where the code is held and continually updated. In the Appendix we provide a computational flowchart, full details of the model parameters and variables, and of extensions based on literature updates. The updates incorporate, among others, cytoplasmic Ca^2+^ and Mg^2+^ buffering, ionophore-mediated transport, the effects of oxy-deoxy transitions on the isoelectric point of haemoglobin, cell pH and [Mg^2+^]_i_ levels, and PIEZO1 channels in the RBC membrane, of particular relevance for the study of RBC homeostasis changes in the circulation in health and disease.

Comprehensive analysis of the full spectrum of changes in RBC homeostatic variables enables here, for the first time, a detailed understanding of the sets of interconnected non-linear processes that modify the intracellular milieu, membrane potential and membrane traffic during K^+^ permeabilization.

### Using the red cell model

The flowchart of figure 1 illustrates the approach applied for the design of the user interface aiming to optimize comparisons between predicted and experimental results, a design guided by the principle that modelling usefulness depends on a close proximity between model and bench. Use of the model will be illustrated in this guide by applying it to an analysis of the “Gardos effect”, trying to harmonize scientific interest with tutorial simplicity.

**Figure 1.**
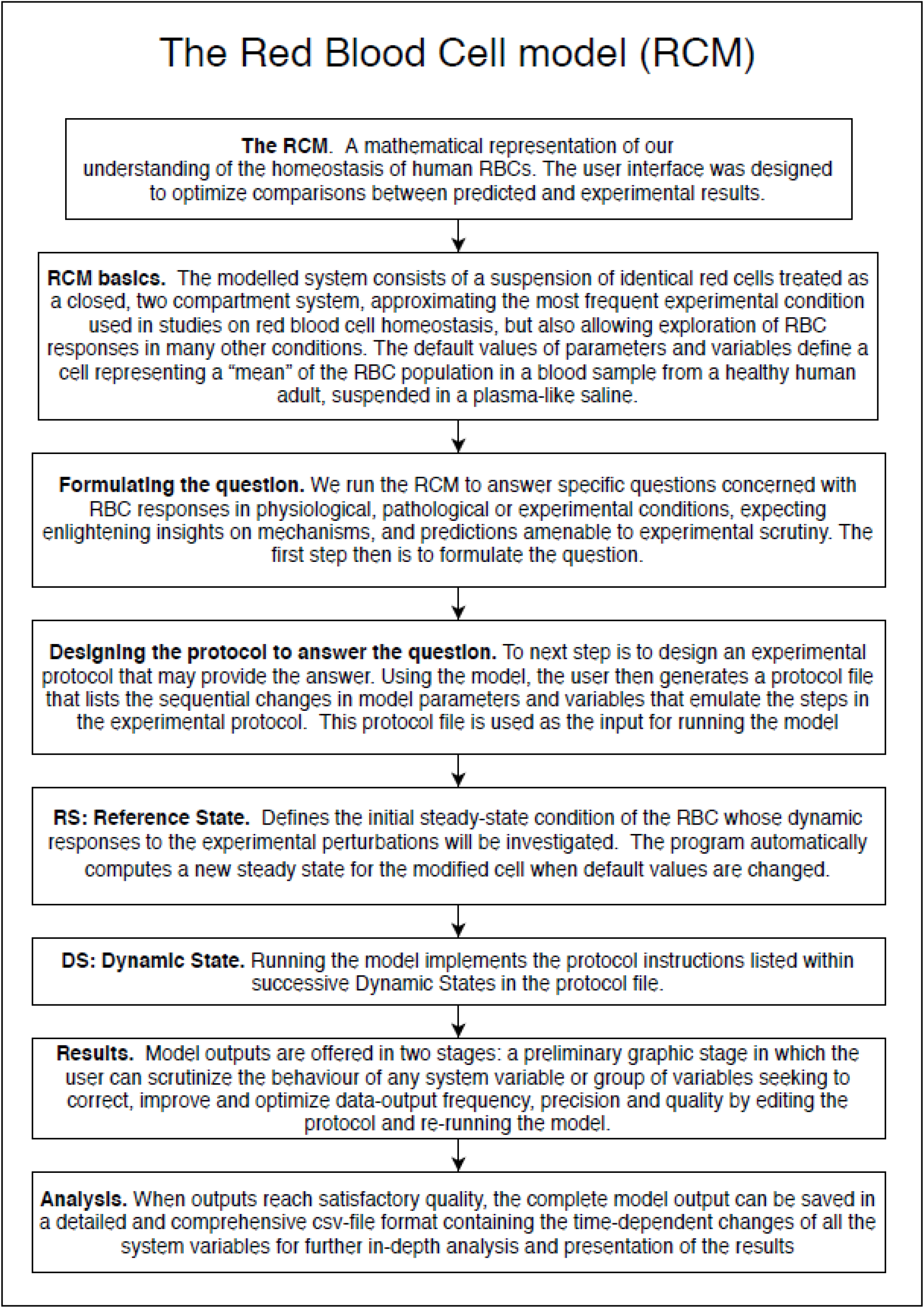
Flowchart illustrating the approach applied in guiding the design of the red blood cell model and the sequential steps involved in running the model.

### Open access to the red cell model

The executable file of the model, RCM*.jar, is available for downloading from the same GitHub repository where this guide is held: https://github.com/sdrogers/redcellmodeljava. The model operates as a RCM*.jar programme within the JAVA environment which needs to be preinstalled. It is recommended not to alter the original file name as it contains coded information on date and update status. Altered names are best applied to shortcuts. Double-click to activate the programme. As explained in the tutorial Boxes below, operation of the RCM model generates protocol files as editable *.txt files, and files containing the results of simulations in spreadsheet format as *.csv files, the columns listing the changes in all the system variables as a function of time. By default, these are saved within the same directory as the one from which the programme is operated, a choice easily modified from within the programme.

### Using the “Gardos effect” to introduce the red cell model (RCM)

Besides its didactic and historic value, the example chosen is of substantial current interest for RBC physiology and pathology. It is based on a puzzling original finding by Gardos [40–42], the “Gardos effect”, a rapidly developing dehydration of red blood cells incubated in plasma-like media following the joint addition of a metabolic inhibitor and a metabolic substrate. Years later, the mystery surrounding the sequential steps involved in the Gardos effect was elucidated by showing that the inhibitor-substrate combinations (iodoacetamide with glucose or inosine, for instance) caused accelerated and profound ATP depletion, the substrate consuming ATP during the initial steps of glycolytic metabolism while downstream ATP production remained blocked by the inhibitor [43–45]. ATP depletion, in turn, reduced Ca^2+^ extrusion through the plasma membrane calcium pump (PMCA) with gradual increases in the concentration of Ca^2+^ in the cytoplasm, [Ca^2+^]_i_ [46, 47]. Elevated [Ca^2+^]_i_ activated a Ca^2+^-sensitive K^+^-selective permeability pathway leading to rapid KCl loss and cell dehydration. Gardos’ landmark discovery of this permeability pathway was eventually recognized as the first report of a calcium-sensitive K^+^ channel, (KCa3.1, KCNN4 gene), baptized as the Gardos channel in the RBC lore [43, 45, 48].

When the Gardos effect was originally simulated [17] the model predicted not only the expected and documented effects of KCl loss and cell dehydration but also some totally unexpected and counterintuitive outcomes, soon experimentally verified [18]. Eventually some of these “side-effects” proved crucial for unravelling the fast track dehydration mechanism of a subpopulation of sickle cells, the irreversible sickle cells [49, 50], whose transient hyperdense state causes widespread vasooclusion and is responsible for most of the clinical symptoms in sickle cell disease [9]. The Gardos channels were considered important participant in the process of progressive RBC densification during circulatory senescence, and channel mutations have recently also been implicated in congenital haemolytic anaemias with altered RBC hydration states [37, 51, 52]. It is the wide scope of the seemingly unrelated homeostatic RBC changes triggered by Gardos channel activation which makes the exploration of the Gardos effect such an excellent didactic choice for introducing the user to the operation of the RCM and for illustrating how the model can be used for an in-depth scientific exploration of the entangled mechanisms behind global cell responses.

Following the sequence outlined in Fig 1, we start by formulating a question and by designing an experimental protocol expected to provide the answer. We simulate that protocol with the model, run the model, report outputs, analyse the results and try to interpret the mechanisms behind predicted responses. Following past experience, the analytical stage often exposes flaws or improvements required in the original protocol leading to further simulations to unravel the complexities of a certain process, in turn guiding the improvements needed in the design of the original experimental protocol, a path illustrated here in the tutorial.

The technical aspects concerned with model operation will be displayed in four separate consecutively numbered text boxes containing detailed instructions on how to construct protocols and run the simulations. Box-separated material will allow readers primarily interested in the science to follow the main text unhindered by details and tutorials related to model use.

#### The protocol-prompting question

With RBCs suspended in plasma-like media at 37°C, what are the effects of sudden and simultaneous inhibition of the Na/K and calcium pumps? With this formulation we bypass the inhibitor-substrate stage of the original Gardos effect and focus on the downstream effects of pump inhibition by ATP depletion. There is also a simple experimental correlate for such a simulation. Vanadate, a well known irreversible inhibitor of P-type ATPases, the family the Na/K and PMCA pumps belong to, instantly blocks pump-mediated transport when added to a RBC suspension, without affecting RBC metabolism [53, 54]. We outline first the vanadate version of the experimental protocol and follow up with the simulated correlate in the model.

#### Outline of experimental protocol

We start with fresh, washed RBCs suspended in a plasma-like saline at a 10% cell volume fraction (0.1 CVF or 10% haematocrit, Hct). The suspension is kept at 37 °C under constant magnetic stirring to prevent cell sedimentation and the formation of significant diffusional gradients. After about 30 min to allow for minor adjustments to the new conditions, we add vanadate to instantly block ion transport through the Na/K and calcium pumps, and follow the evolution in time of system variables, the vanadate version of the Gardos Effect protocol.

#### Outline of simulated protocol

In the simulations we bypass the usual preparatory steps from drawing blood to washing the RBCs free of plasma, white cells and platelets. We start with a 0.1 cell volume fraction (CVF, equivalent to a 10% Hct) in a plasma-like, isosmotic HEPES-Na-buffered saline solution, and probe system stability while incubated at 37°C over a control period. Vanadate addition is represented by sudden 100% inhibition of Na/K and PMCA pumps, and the model follows the evolution in time of all the variables in the system. Because the cell-medium system is treated as a closed, two-compartment system, mass conservation applies, and redistribution equations compute the composition changes in each compartment at each instant of time. Instant, gradient-free uniformity in the composition of cell and medium compartments is assumed throughout representing the approximation intended by magnetic stirring in the experimental protocol. Simulated protocols are stored in text files, and model outputs reporting the changes in all the homeostasis variables of the system as a function of time are saved as comma separated (csv) spreadsheets.

The instructions for entering protocol simulations are contained in two separate sections in the model: the Reference State (RS) and the Dynamic State (DS). The RS defines the initial constitutive condition of the RBC under investigation. In the DS section the user enters the protocol prescribed sequence of perturbations to parameters and variables in consecutive DS pages, automatically numbered by the programme. All user entries in RS and DS modify default-suggested values. Text windows within each of the RS and DS pages provide contextual guiding information. Separate RS and DS Help pages provide additional information for users to consult.

In Boxes 1-4 next, the reader is guided step by step on how to simulate the vanadate version of the Gardos effect protocol using the RCM*.jar executable. Although the tutorial remains focused on the chosen example, it is hoped the user will note, along the way, the vast range of experimental simulations the software enables within a rather austere display of choices. The model output, stored in csv format, contains the evolution in time of all the homeostatic variables considered in the model during the successive protocol-defined stages for subsequent plotting and analysis. In the Results and Discussion sections we illustrate how this data can be used to report the results and to explore the mechanisms behind each of the predicted responses.

###### Box 1: The Reference State

Figure 2 shows the first dialog box that comes up when we activate RCM.jar. It contains a Welcome message and two optional prompts at the bottom. Select New Experiment, as there are no yet stored protocols on file. This brings up the central page for simulating experiments (Fig 3). This page contains two main panels and five tags at the bottom. The left panel, the Reference State (RS), defines the initial constitutive condition of the RBCs at the start of experiments. On the right panel, the Dynamic State (DS), the user specifies the duration of the experiment, the frequency and precision of the data points with which results will be reported, and the changes in the value of selected parameters and variables meant to represent the successive perturbations the RBCs are exposed to in the course of experiments. Press the HELP tag at the bottom for a brief guide to all entries in the RS and DS panels, and to those in the dedicated DS PIEZO tag. The default values assigned to each of the parameters and variables in the Reference State and in the Dynamic State, listed within each of the tags, represent rounded *mean values* obtained from the literature for RBC samples from healthy adults (see Appendix for glossaries). It is important to note that when using the RS default values, model predictions will be reporting the responses of this particular *mean* representative cell. With a mean circulatory lifespan of about 120 days, real RBC samples are a mix of about 120-day cohorts. Because many constitutive variables change with RBC age it is important to bear in mind modifying the RS default values when exploring responses associated with RBC age diversity. For the Gardos effect simulation we will accept the default Reference State values of the “mean” RBC and proceed to simulate the successive experimental steps within the Dynamic State right panel (Fig 2).

**Figure 2.**
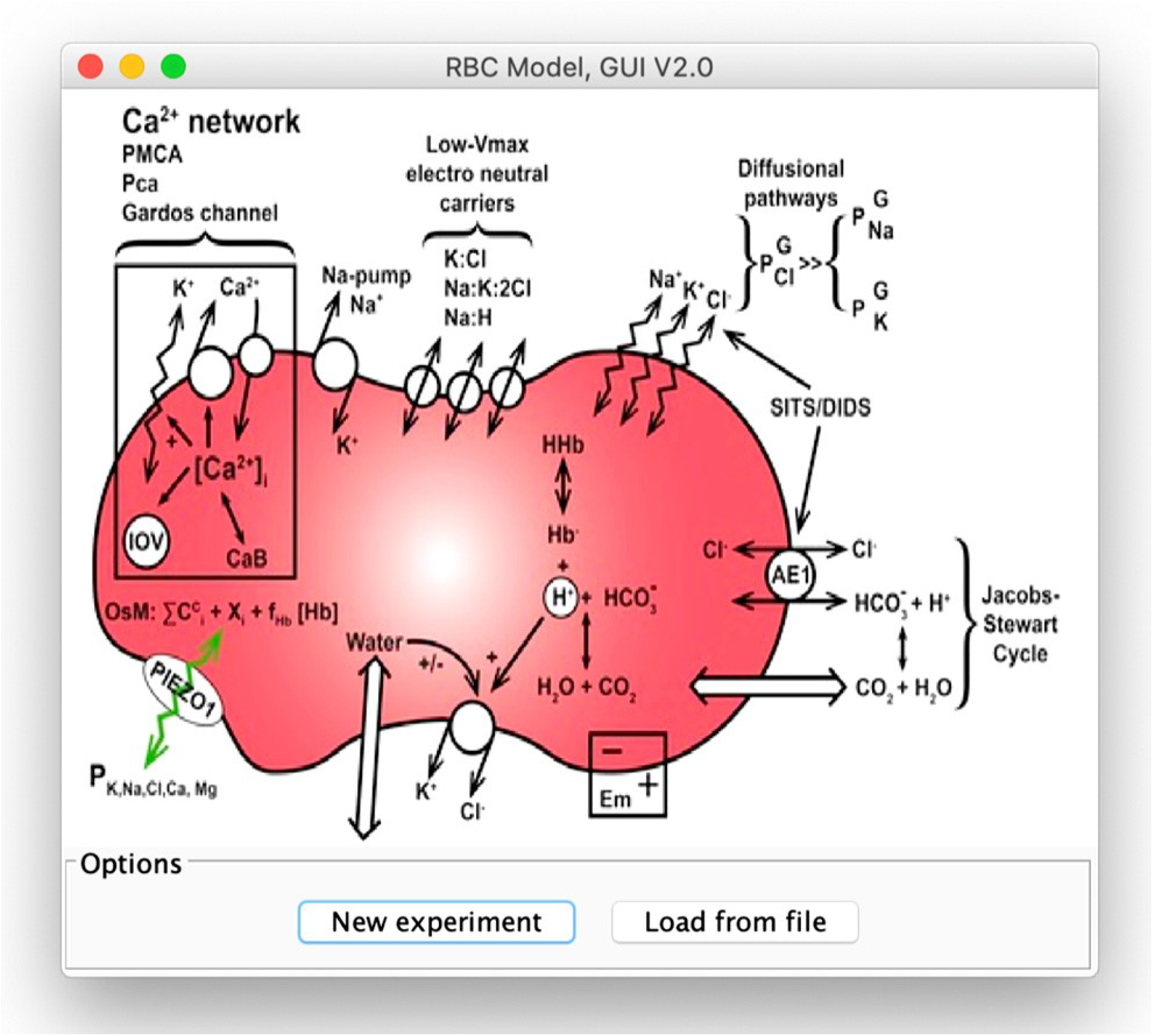
The Welcome Page. The figure contains a simplified representation of the main components of the RCM, including PIEZO1. At the bottom of the page the user is offered two tags with alternative options to create a new simulated protocol, “New experiment” (on the left), or to transfer a previously saved protocol file “Load from file” (on the right).

**Figure 3.**
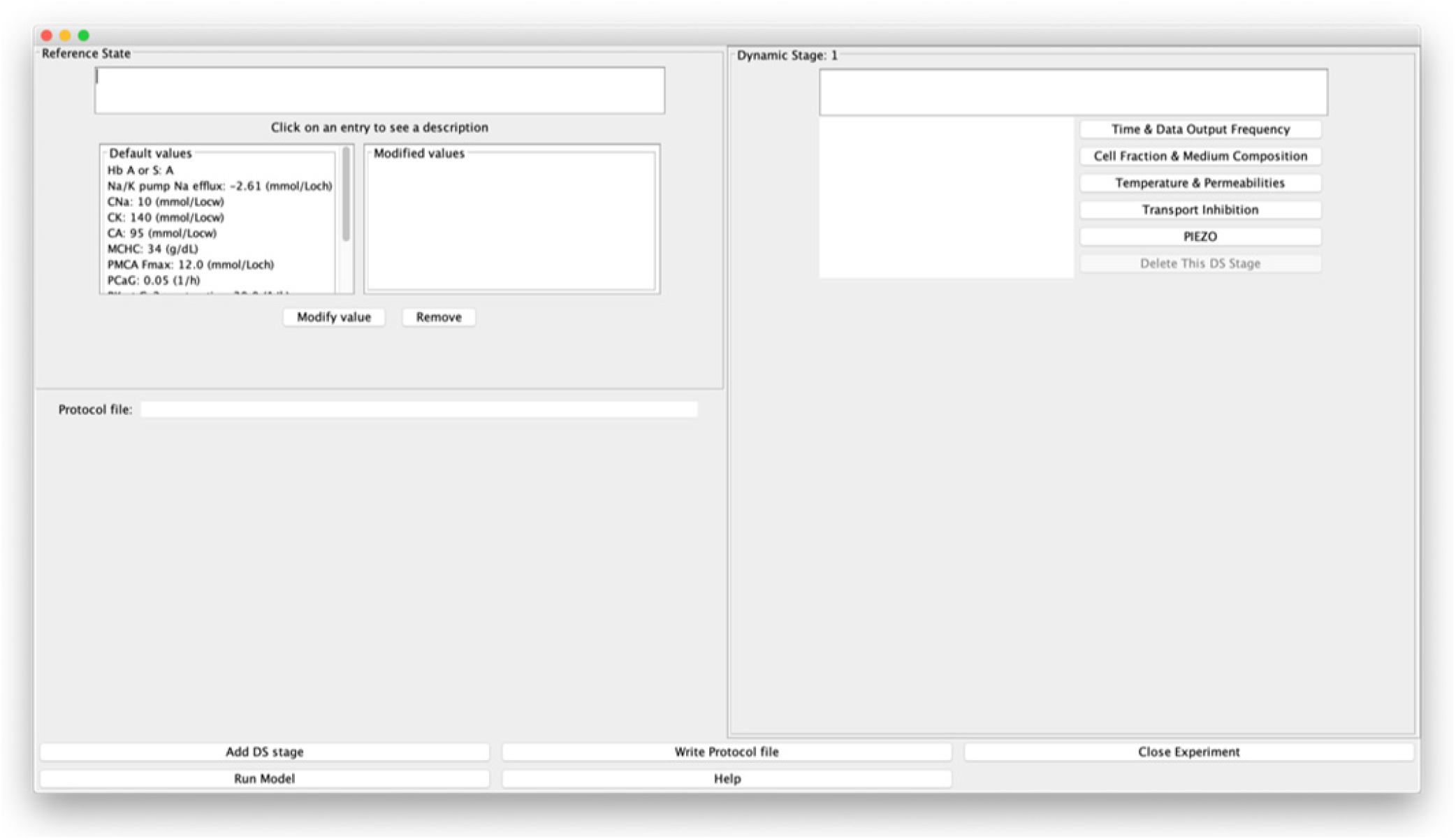
The Central Page (reduced version), with options for changing the initial (default) constitutive properties of the cell in the Reference State (RS, left panel), and to implement sequential Dynamic States simulating experimental steps (DS, right panel). Detailed operation is explained in boxes 1-4 in the text using an example of a simulated experiment.

###### Box 2: The Dynamic States

The outlined protocol requires implementation in two DS stages, DS1 and DS2, the first to allow the system to settle into whichever new conditions are set and provide baseline values of variables before the next experimental stage, the second to simulate the Vanadate-version of the Gardos effect with sudden inhibition of the Na/K and calcium pumps. Start the DS1 stage by selecting the “Time & Data output frequency” tab on the right. This opens up a full dialog box on the left. You select “time” and then “Modify value”, or simply double click on “time” to open the “Input” window where you enter the new or modified value; let us use 30 min in this instance. Note how the “time 30” instruction becomes registered on the left-side window of the DS1 panel. Errors are easily corrected by repeating these steps with amended entries. Now select the “Cell fraction & medium composition” tab. Operating the prompts in the way instructed above for “time” proceed to enter 0.1 for the cell volume fraction (CVF) to represent a 10% haematocrit. Leave all other variables unchanged, as the default medium composition approximates widely used conditions for “plasma-like”, protein-free media used in experiments with RBCs. Note how the changes become incorporated in the protocol window of DS1.

Next click on “Add DS stage” to add a DS2 stage. Set time to 60 min and enter 100 in the Transport Inhibition tab for 100% inhibition of both the PMCA and the Na/K pump. The cell fraction remains as set for DS1. *Instructions are automatically carried over sequential DS stages unless specifically deleted or changed*.

###### Box 3. Running the model, inspecting preliminary results, and considering the desirability of protocol improvements

We are now ready to run the simulation. Good practise recommends saving the protocol at this stage, for immediate availability if changes become necessary. Press the “Write a protocol file” tag at the bottom of the central page (Fig 3). Choose a directory and filename (GE1 for instance). The protocol is saved as a text-file with name GE1.txt, editable in any text editor. Figure 4A shows what the text-file looks like at this stage. Although it is possible for the user to modify entries in the text file, we envisage the file mainly being used for archiving and allowing protocols to be saved and reloaded later. To minimise the chance of user error, we recommend all protocol changes to be made from within the central page of the user interface.

**Figure 4.**
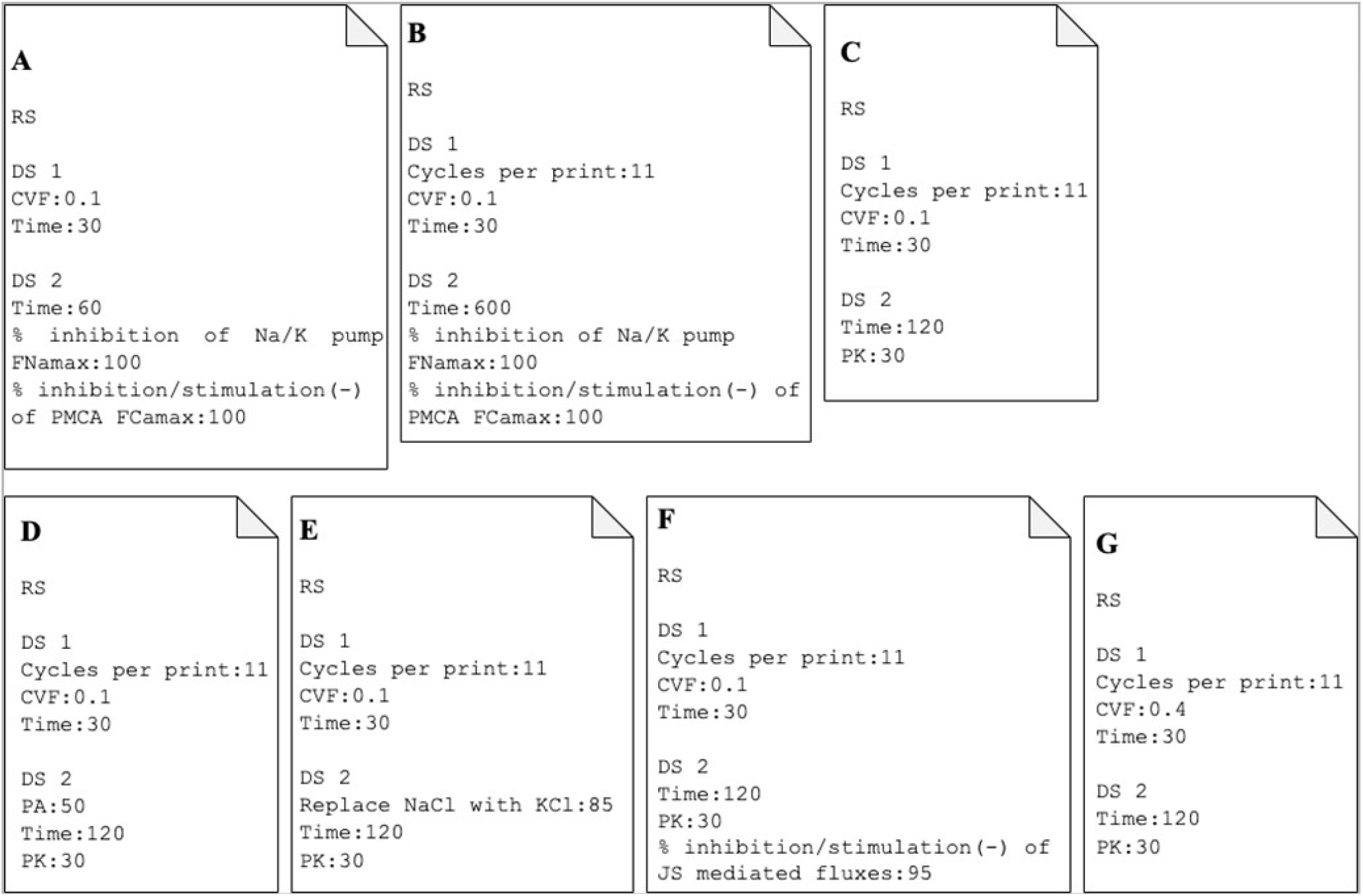
Alternative protocols. Samples of protocol text files on the K^+^-permeabilization theme. Users are advised to save result files (csv) for each of these protocols to practise interpreting the mechanisms responsible for the different responses, with particular attention to the following variables: RCV, Em, pHi, rA-rH, COs-MOs, CK, CNa, CA. **A** and **B**: show the original and corrected protocol versions outlined in boxes 3 and 4 of the tutorial, respectively. **A** and **B** are the only protocols explicitly analysed in the text. **C: Effect of valinomycin**. Addition of the K^+^-selective ionophore valinomycin bypasses all calcium-mediated effects. This basic protocol is repeated with variations in all the following examples (D to G). Note the rapid onset of hyperpolarization and dehydration caused by sudden K^+^ permeabilization, relative to their slower onset rates under the vanadate-simulated Gardos effect (shown in Figs 5A and 5C). **D: Increased anion permeability (PA in DS2)**. Comparison of dehydration rates between protocols **C** and **D** illustrates the powerful rate-limiting effect of the anion permeability on RBC dehydration rate; the electrodiffusional anion permeability can be increased experimentally by replacing 10 mM NaCl with 10 mM NaSCN in the medium [55]. **E: Iso-osmotic replacement of 85 mM NaCl for KCl in the medium**. Elevating the medium potassium concentration effectively reduces the driving force for dehydration and for all associated secondary effects. **F: Effects of AE1 inhibitors**. Dehydration remains rapid and profound, but the secondary JS-mediated pH effects are retarded, with rH approaching rA slowly instead of almost instantly as for the un-inhibited AE1 exchanger in the Gardos effect simulations (Fig 8D). **G: Osmotic effects of RBC dehydration explored at high haematocrits**. The cell fraction is set to 0.4 (40% haematocrit) to illustrate how the osmotic effects of haemoglobin crowding become more marked the higher the cell fraction and overall haemoglobin content of the suspension. Compare the COs and MOs variables in conditions **C** (with 0.1 cell volume fraction) and **G**.

**Figure 5:**
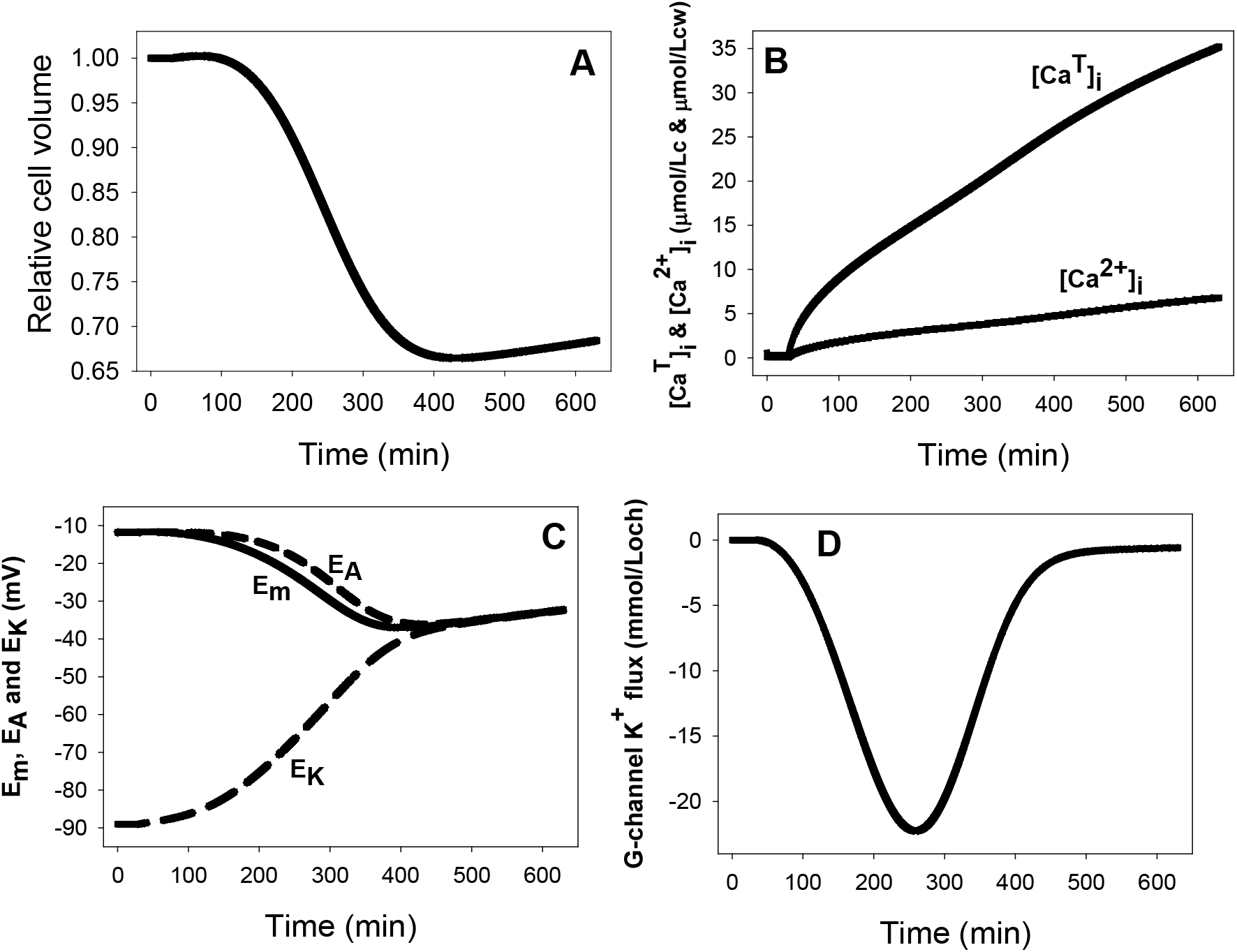
Predicted changes in selected RBC variables as a function of time following sudden inhibition of the Na/K and calcium pumps by vanadate. **A:** relative cell volume changes due to Na/K pump inhibition and to delayed activation of Gardos channels by slow [Ca^2+^]_i_ gain; **B:** total ([Ca_T_]_i_) and free ([Ca^2+^]_i_) intracellular calcium concentration changes following PMCA inhibition; **C:** Changes in membrane potential, E_m_, and in the equilibrium potentials of K^+^ and A^−^, E_K_ and E_A_, respectively, with time-patterns determined by Gardos channel activation kinetics and by ion gradient dissipation; **D:** biphasic net K^+^ flux pattern determined by changes in Gardos channel permeability (increasing K^+^ efflux phase) and K^+^ gradient dissipation (decreasing K^+^ efflux phase), with minimal initial and late contributions of Na/K pump inhibition, hardly noticeable on this y-axis scale.

Select “Run Model” at the bottom of the central page. This opens a window on the left overlapping the central page and showing the progress of the model computations, almost instant for this protocol. At the bottom, four commands offer data saving, window closing, stopping model run, and data plotting options. The “Stop” command halts the model run; useful when exploring protocol conditions to optimize the compromise between data output precision and overall duration of run. The data plotting option is particularly useful for a preliminary exploration of results before saving data, to enable protocol adjustments on the go before reaching worth-saving outputs. Let us illustrate this by reviewing the data outputs from the current protocol.

Press the Launch plotter command. This opens a small window with a comprehensive list of all the model variables. Our interest is directed primarily at the variables most influenced by PMCA inhibition and Gardos channel activation: cell volume, membrane potential, potassium flux and cell calcium. Press the relative volume variable, V/V. This brings up a basic graph of V/V as a function time for the set duration of the experiment. Scaling of the y-axis automatically adapts to the magnitude of change in the selected variable. Within the first hour after pump inhibition, we note a few points outlining a tiny increase in cell volume followed by an accelerating decrease. But with such poor quality of data it is impossible to derive any significant information. It is clear from this graph that we need a much higher frequency of data output points and a much more extended time-scale to explore properly the predicted effects.

###### Box 4. Correcting the protocol using graphic output feedback and saving improved final model output

Let us return to the original protocol to increase data output frequency and experiment duration. Close any open plot windows, and the window showing model progress. In DS1 press the “Time & Data output frequency” tab and change the “cyclesperprint” from its default value of 777 to a lower value, 11 in our example (thus forcing the model to output values at a much higher frequency). In DS2 change “time 60” to “time 600” to explore effects over a longer period. Save the modified protocol overwriting the original one using the same name (Fig 4B). Now run the model again. Note that this time the model run takes longer in line with the increased density of output data and duration of the experiment. Launch the Plotter and explore the time changes of the variables listed as of primary interest: V/V, QCa, CCa2+, Em, and FKGardos. Note the smooth appearance of the curves with the new setting of “cyclesperprint” at 11.

The increased data-sampling frequency and duration of the experiment seem to adequately cover now the period of major changes and deliver smooth data outputs, rendering outputs well-suited for an in-depth analysis of the full RBC response to the vanadate-simulated Gardos effect. To save the full complement of data generated by the model press the Save Output tab and at the filename prompt enter GE1. This generates a file named GE1.csv open to scrutiny and graphing in different software platforms. We will now generate figures from this file. These should guide the user on how to report the results, and on how to analyse and interpret the data in search of the mechanisms behind the observed effects.

## Results and Analysis

### Predicted time-course of changes in RBC volume, membrane potentials, calcium contents and Gardos channel mediated K+ flux following the vanadate-protocol simulation of the Gardos effect

Following pump-inhibition, the change in RCV (Fig 5A) shows three distinct phases, a slow initial increase, a sharp decline followed by a shallow slow recovery. The initial and late slow rates of volume increase result from Na/K pump inhibition, as may be shown by re-running the vanadate protocol without Na/K pump inhibition. The reason for such slow rates of volume change and cation gradient dissipation is the extremely low permeability of the RBC plasma membrane to Na^+^, K^+^, and to cations in general, an evolutionary optimized condition allowing RBC volume maintenance with minimal pump-leak turnover rates and energy consumption [6].

The sharp volume fall during the second phase results from Gardos channel activation (Fig 5D). The delay is caused by the slow increase in [Ca_T_]_i_ and [Ca^2+^]_i_ following PMCA inhibition (Fig 5B). The low Ca^2+^ permeability of the RBC membrane limits the rate at which [Ca_T_]_i_ increases, and cytoplasmic Ca^2+^ buffering further reduces the scale of the [Ca^2+^]_i_ increase which is the direct signal for Gardos channel activation. Figures 5C and 5D show the time-course associated with hyperpolarization and channel-mediated K^+^ flux, respectively. It can be seen (Fig 5C) that the membrane potential follows closely the anion equilibrium potential throughout all the periods, reflecting the dominance of anion conductance in the RBC membrane [56–58]. The E_m_-E_A_-E_K_ convergence in the end (Fig 5C) reflects the new ion-gradient distribution generated after full dissipation of the K^+^ gradient. The K^+^ efflux curve through the Gardos channel (Fig 5D) shows a biphasic pattern, first a Ca^2+^-activated increase followed by a decline as the K^+^ concentration gradient dissipates.

Interested readers may wish to explore further and compare the Gardos effect response described here with that elicited by sudden K^+^ permeabilization of RBCs. Sudden K^+^ permeabilization can be elicited experimentally by addition of valinomycin [18, 59], a potassium-selective ionophore, as simulated with the protocols of Figs 4C to 4G, or by addition of a Ca^2+^ ionophore [60–62]. Note the sharp differences in the rate and magnitude of changes in membrane potential and other variables between different K^+^-permeabilization protocols.

### Predicted time-course of changes in RBC pH and system osmolarity following K^+^ permeabilization

A first set of predictions, totally unsuspected when originally advanced [17], concerned changes in cell and medium pH, with cell acidification and medium alkalinisation (Fig 6A), predictions experimentally validated in a variety of experimental conditions [18, 28]. Their mechanism is explained in the Discussion. The biphasic time-course predicted for the net H^+^ flux in this protocol is shown in Fig 6B. Further scrutiny of csv-file data shows that net K^+^ flux exceeds net Cl^−^ efflux, and that flux electroneutrality is made up by H^+^ influx, with the appearance of an electroneutral K^+^:H^+^ exchange, misattributed to a real electroneutral K^+^:H^+^ antiport in the past [63–65]. Its phantom nature was exposed by the original model which, lacking this transporter, predicted most of the observed experimental results used in its support and explained the complex mechanisms behind its appearance [17]. The non-existing K^+^:H^+^ antiport carries instructive cautionary advice on applying a strict minimalistic approach when building up model components beyond the essential basic components.

**Figure 6.**
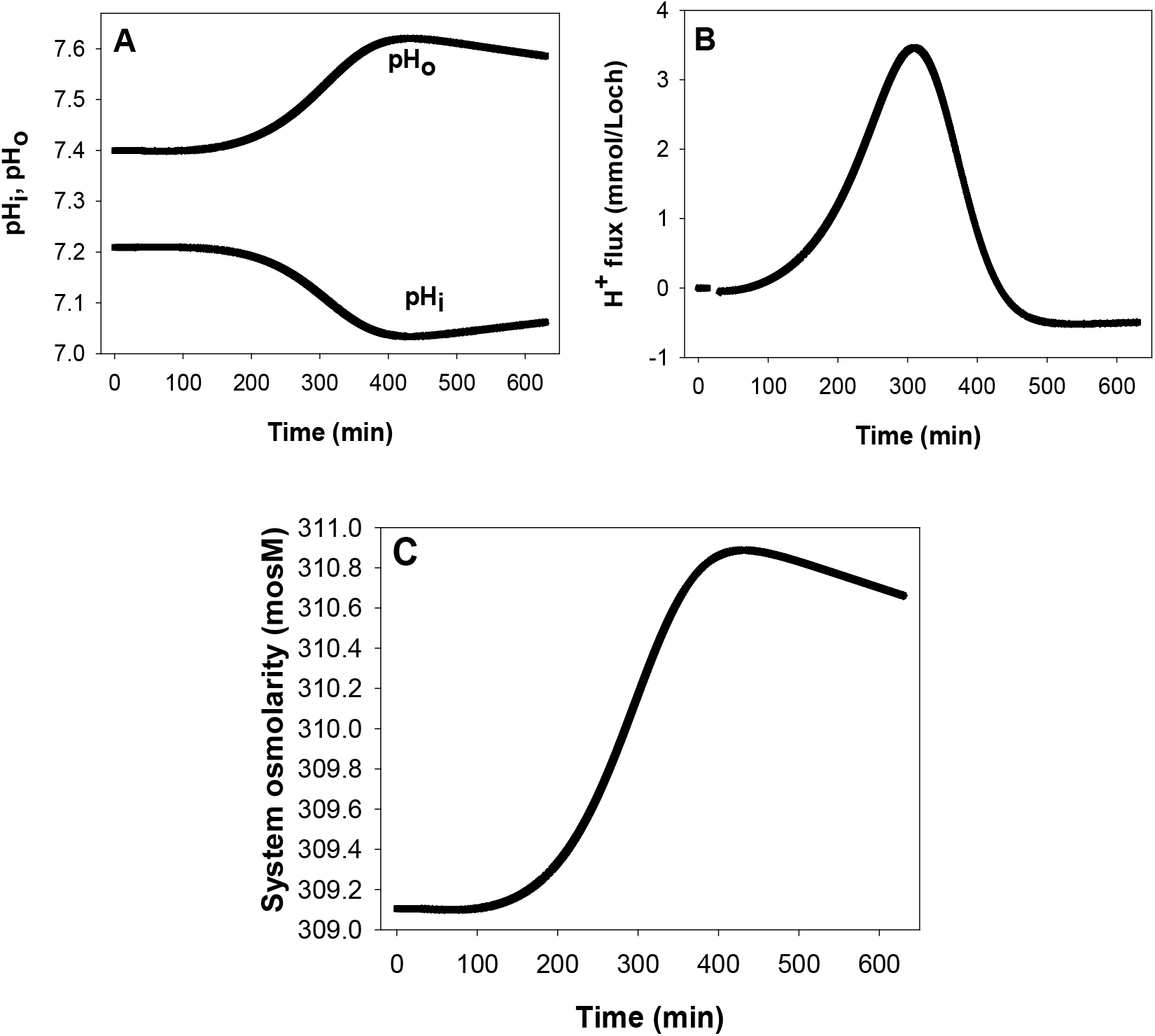
Predicted changes in pH and system osmolarity following the vanadate protocol. **A:** Changes in cell and medium pH (pHi and pHo, respectively) with patterns determined by the changes in Cl^−^ concentration ratio, rA, across the red cell membrane; **B:** biphasic H^+^ flux pattern through the JS mechanism driven by the changes in rA; **C:** changes in system osmolarity resulting from variations in haemoglobin crowding secondary to cell volume changes.

A second set of unexpected predictions concerned system osmolarity (Fig 6C). The model predicts that cell dehydration driven by net KCl loss elevates cell and medium osmolarity as if new osmotic particles were generated within the cells and redistributed between cells and medium. Moreover, the model predicts that the increase in global system osmolarity will be higher the higher the cell fraction, an effect the reader is invited to explore further (Fig 4G).

## Discussion

We introduced a multiplatform JAVA-based model of human red blood cell homeostasis and applied it here to explore in detail the homeostatic changes induced by activation of Gardos channels while guiding users on how to operate the model.

The deceptive simplicity of the central page (Fig 3) for determining the constitutive properties of the cell in the initial Reference State, and for the design and creation of simulated experimental protocols in the Dynamic State hides an unlimited versatility in the range and scope of questions that can be addressed with the model. Availability of the continually updated code in the online GitHub repository also allows users to extend the model for specific uses not included in current versions. We hope the accessibility and simplicity of RCM*.jar will encourage its use as a frequently consulted tool on questions related to RBC volume, transport and homeostasis in health and disease.

We discuss next the mechanisms concerned with pH and osmolarity changes associated with K^+^ permeabilization, good examples of entanglement confounding intuition.

### Predicted side-effects of K^+^ permeabilization of human RBCs

Let us consider first how the operation of what has become known as the Jacobs-Stewart cycle or mechanism [66, 67] is represented in the model. The JS cycle comprises the parallel operation of the anion exchanger, AE1, as an electroneutral Cl^−^:HCO_3_^−^ antiport, and the CO_2_ shunt, as illustrated in Fig7A.

**Figure 7.**
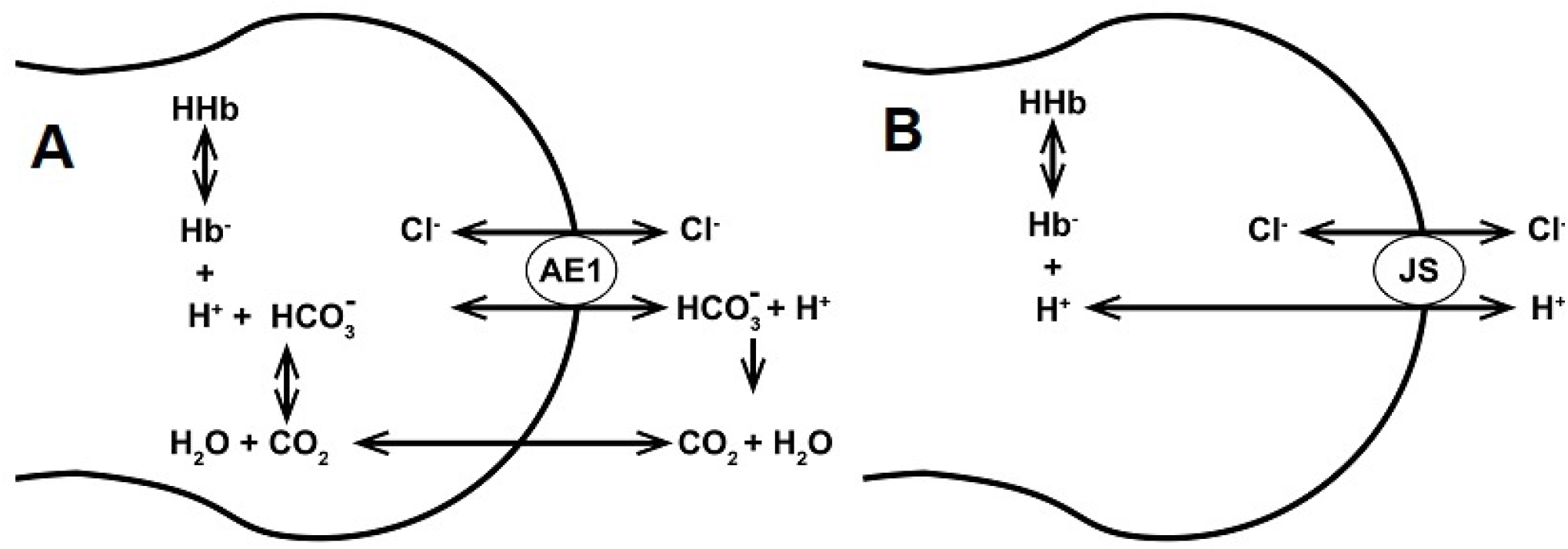
Representation of the JS cycle as an electroneutral Cl^−^:H^+^ symport. **A:** The Jacobs-Stewart mechanism showing the combined operation of the anion exchanger, AE1, and the CO_2_ shunt on the transfer of H^+^ ions across the red cell membrane; **B:** equivalent model-representation of H^+^ transport through the JS mechanism as an electroneutral Cl^−^:H^+^ cotransport.

As a rough approximation to a complex multistep process, we may assume that for each Cl^−^:HCO_3_^−^ exchanged through AE1, protonation of the transported HCO_3_^−^ ion effectively adds a CO_2_ molecule to the medium on the membrane side it was transported to. Rapid CO_2_ equilibration across the membrane shunts back the CO_2_ to the other membrane side releasing a H^+^ on hydration. The net result of each half JS cycle is therefore the electroneutral transfer of one Cl^−^ with one H^+^, indistinguishable from the operation of an electroneutral Cl^−^:H^+^ symport. Thus, the complete JS cycle may be represented phenomenologically by an electroneutral Cl^−^:H^+^ cotransporter, as illustrated in Fig 7B, ignoring the background roles of HCO_3_^−^ and CO_2_. This phenomenology lumps a large number of steps operating over a vast range of time-scales comprising CO_2_ solvation, CO_2_ hydration (spontaneous in the extracellular medium and carbonic anhydrase-assisted intracellularly), CO_2_ transport, intracellular and extracellular proton buffering, and the complex kinetics of the AE1 exchanger. The Cl^−^:H^+^ cotransport representation has been shown to accurately describe the operation of the JS cycle at the functional level required for modelling RBC homeostasis. It allows the model to predict pH changes and explain the mechanisms behind all relevant physiological, pathological and experimental RBC responses involving the participation of the JS cycle [12, 18, 20, 25, 26, 28, 68–70]. An additional important issue addressed by this phenomenology is that of the “ignored” HCO_3_^−^ and CO_2_ (see Box 5). This oversight, together with the JS cycle phenomenology, also enables the model to be run using a single diffusible anion representation, A^−^, for the combined contents or concentrations of Cl^−^ and HCO_3_^−^.

##### Box 5. The ignored HCO_3_^−^ and CO_2_

In most of the experiments with RBC suspensions HCO_3_^−^ salts are not added to the medium and both HCO_3_^−^ and CO_2_ remain unreported as explicit components of the system. Where then is the HCO_3_^−^ that keeps the Cl^−^:HCO_3_^−^ exchange in motion and allows the Cl^−^:H^+^ symport phenomenology to represent the operation of the JS system? Dissolved HCO_3_^−^ at equilibrium with atmospheric CO_2_ is always present in unsealed cell suspensions, and this presence is sufficient to sustain near physiological rates of JS cycle turnover. It can easily be estimated that at an atmospheric pressure of 760 mmHg, a ~ 0.04% CO_2_ in environmental air, a solubility constant of CO_2_ in water of 0.03mM/mmHg, a pKa of 6.147 for the [H^+^][HCO_3_^−^]/[CO_2_] hydration reaction, and at a medium pH of 7.4-7.5, the unreported HCO_3_^−^ concentration will be around 200 μM, more than sufficient to sustain high rates of JS-mediated fluxes.

With over 10^6^ AE1 units per cell and essentially instant CO_2_ equilibration rates across the RBC plasma membrane, the JS mechanism, optimized by evolution for effective CO_2_ transport from tissues to lungs, is by far the most important mediator of H^+^ transport in RBCs. The JS phenomenology is represented in the model by a low-saturation equation of the form F_ClJS_ = F_HJS_ = k_AE1_([Cl^−^]_o_[H^+^]_o_ – [Cl^−^]_i_[H^+^]_i_), where F_ClJS_ and F_HJS_ are the net fluxes of Cl^−^ and H^+^ through the JS pathway, and k_AE1_ is the rate constant of AE1 turnover, about four to six orders of magnitude faster than that of any other ion transporter in the RBC membrane. In the absence of net JS fluxes, F_JS_ = 0, [Cl^−^]_o_[H^+^]_o_ = [Cl^−^]_i_[H^+^]_i_. Defining the concentration ratios rA and rH by rA = [Cl^−^]_o_/[Cl^−^]_i_ and rH = [H^+^]_i_/[H^+^]_o_, respectively, the equilibrium condition for the JS system at zero net flux will be rA = rH. The participation of other anion and proton transporters causes deviations from this equality, miniscule in normal steady states. Larger deviations during dynamic states would tend to be speedily restored towards rH ≈ rA by the JS system.

Let us now consider how RBC dehydration caused by K^+^ permeabilization affects cell pH and osmolarity from the data stored in GE1.csv, the output file generated from the vanadate protocol simulations (***Box 4***). As the cells start to dehydrate by the net loss of KCl, four confluent changes among many others are particularly relevant for the pH and osmolarity effects: i) the concentration of the impermeant intracellular anion increases (Fig 8A, [X^−^]); ii) the concentration of haemoglobin increases (Fig 8B, [Hb]) and so does its contribution to the net negative intracellular charge (Fig 8B, n_Hb_[Hb]); iii) the osmotic coefficient of haemoglobin increases along a power function of the haemoglobin concentration increasing sharply the osmotic contribution of haemoglobin as dehydration proceeds (Fig 8C), and iv), the intracellular concentration of Cl^−^ becomes markedly reduced (Fig 8A, [Cl^−^]) by net Cl^−^ loss, by charge displacement from increasingly concentrated impermeant anions X^−^ and Hb^−^, and by water retention due to the increasing colloidosmotic pressure on haemoglobin.

**Figure 8.**
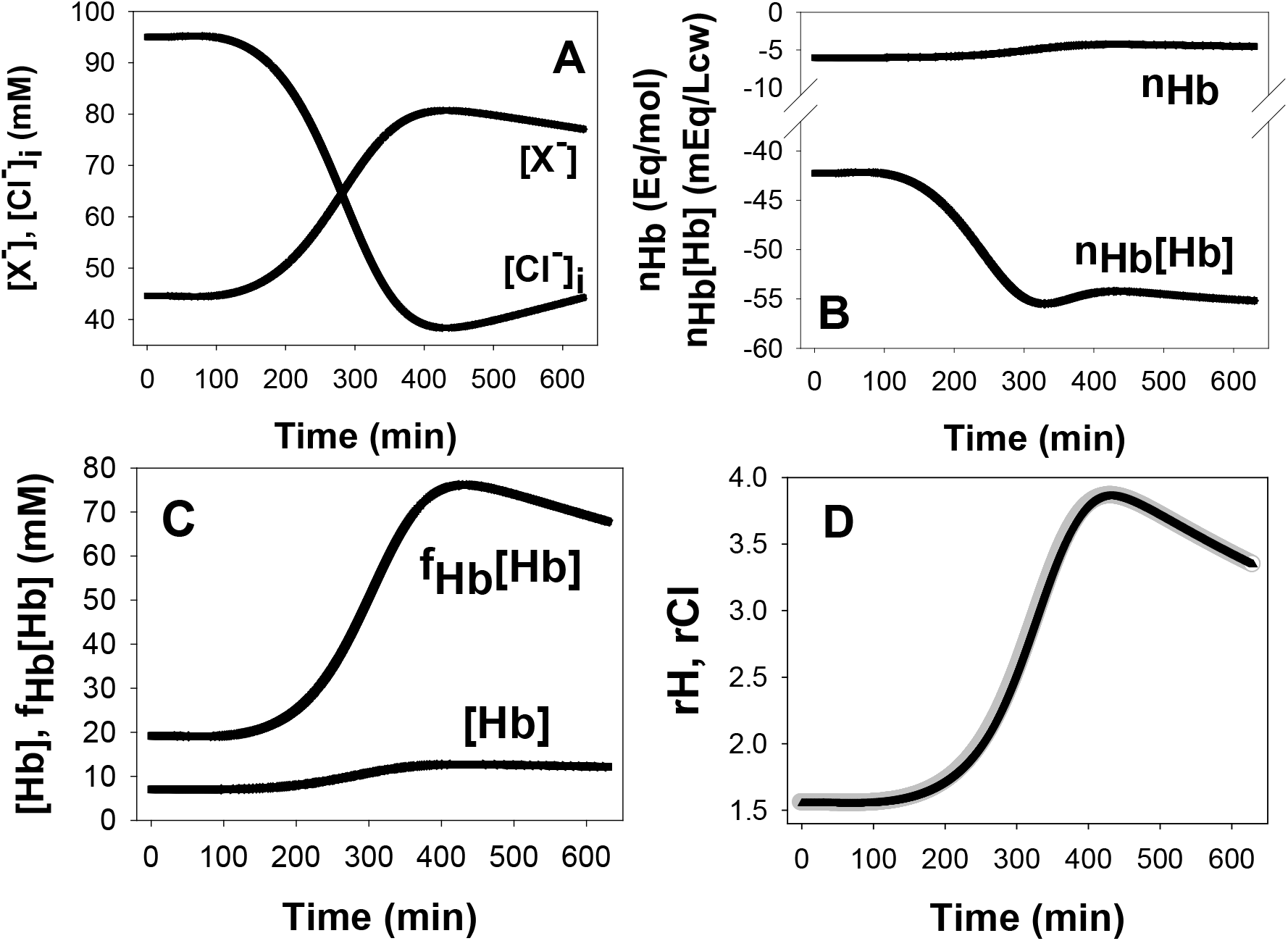
Predicted effects of macromolecular concentration changes on cell Cl^−^, H^+^ and osmolarity. **A:** Inverse correlation between the changes in the concentrations of cell permeant A^−^ ((Cl^−^), [Cl^−^]_i_) and of impermeant cell anions ([X^−^]) during the Gardos effect. **B**: opposite changes in the charge per mole of haemoglobin (n_Hb_) and in the charge concentration on haemoglobin (n_Hb_[Hb]) during the Gardos effect. **C:** illustrates the powerful effect of the osmotic coefficient of haemoglobin (f_Hb_) in elevating the osmotic contribution of haemoglobin during concentrative haemoglobin crowding. **D:** overlapping changes in rA^−^ (gray) and rH^+^ (black) during the Gardos effect highlighting the almost instant equilibration of rH^+^ to rA^−^.

The reduction in [Cl^−^] is the most relevant drive for the pH changes. The resulting increase in rA will tend to be rapidly equilibrated towards rH ≈ rA through the JS mechanism (Fig 8D), explaining the pH changes shown in Fig 6A. Although cell acidification is buffered by haemoglobin with reduction in its net negative charge per mol, n_Hb_ (Fig 8B, n_Hb_), the increase in haemoglobin concentration dominates, and its overall contribution to the intracellular negative charge increases with dehydration (Fig 8B, n_Hb_[Hb]). Electroneutrality is strictly preserved through all these changes, ([Na^+^]_i_ + [K^+^]_i_ + 2[Mg^2+^]_i_ = [A^−^]_i_ + [X^−^] + n_Hb_[Hb]), and the extra osmotic load generated by haemoglobin crowding (Fig 8C) is rapidly redistributed between cells and medium by an hypertonic KCl + KOH effluent [18]. In Box 6 we provide a highly simplified example to illustrate the dehydration-induced acidification mechanism bypassing the complexities generated by macromolecular crowding and proton buffering.

##### Box 6. Thought experiment to illustrate how dehydration by KCl loss acidifies cells that express the anion exchanger (AE1) in their plasma membrane

We imagine a protein-free RBC ghost resealed with 50 mM KGluconate, 100 mM KCl, and 1 mM ATP, suspended in a 150 mM NaCl medium adjusted to a pH_o_ of 7.4, equivalent to a [H^+^]_o_ of ~ 40nM. Monovalent gluconate is the only impermeant solute in this system. ATP is there to fuel the Na/K pump and balance the passive Na and K leaks to set ghosts initially in a steady state. We add valinomycin to permeabilize the ghost to potassium eliciting its dehydration by potassium gradient-driven loss of an isosmotic KCl effluent. We wait until the ghost dehydrates to half its initial volume (V/2) and take note of the change in its composition.

| Cell concentrations (mM) | Initially | After $V/2$ |
| --- | --- | --- |
| Gluconate | 50 | 100 |
| $\text{Cl}^-$ | 100 | 50 |
| $\text{K}^+$ | 150 | 150 |

With rA defined by rA = [Cl^−^]_o_/[Cl^−^]_i_ and rH by rH = [H^+^]_i_/[H^+^]_o_, the JS-AE1 mechanism sets rH to match rA at a rate determined by AE1 expression and turnover. Therefore, when rH ≈ rA,

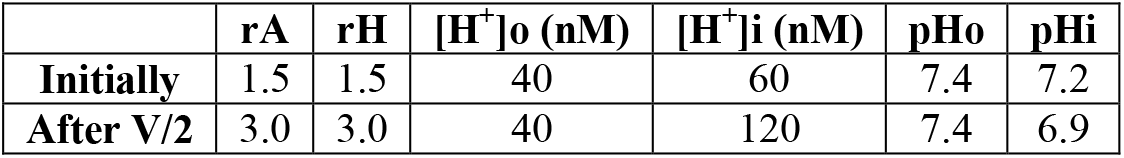

| | $r\text{A}$ | $r\text{H}$ | $[\text{H}^+]_o$ (nM) | $[\text{H}^+]_i$ (nM) | $\text{pH}_o$ | $\text{pH}_i$ |
| --- | --- | --- | --- | --- | --- | --- |
| Initially | 1.5 | 1.5 | 40 | 60 | 7.4 | 7.2 |
| After $V/2$ | 3.0 | 3.0 | 40 | 120 | 7.4 | 6.9 |

The dehydration-induced increase in rA led to a fall in intracellular pH from 7.2 to 6.9 in this example. Cell acidification following dehydration led by chloride salt losses has been documented in cells other than RBCs a tendency common to any cell type expressing the AE1 in its plasma membrane [65].

Haemoglobin crowding (Fig 6C) has three main side-effects: i) it contributes to increase the net negative charge on impermeant cell anions (Fig 8B), ii) it elevates cell and system osmolarity (and osmolality) as a power-function of its concentration (Figs 6C and 8C), and iii) it increases the colloidosmotic pressure within the cells causing water retention and a progressively more hypertonic effluent as part of the redistribution process of permeant ions between cells and medium (Fig 6C). It tempts speculation that this powerful osmotic effect of soluble protein crowding, so easily detectable in a RBC suspensions, may yet be discovered to participate in the fluid dynamics of cells containing compartments with a high concentration of soluble macromolecules.

We can now summarize the sequence of events during the Gardos Effect within the scope of conditions considered in this Gide (Fig 9). Running down of cell ATP by the combination of substrate and inhibitor reduces Ca^2+^ extrusion by the plasma membrane calcium pump. This allows a net build up of the intracellular concentration of free calcium to levels that activate Gardos channels (Figs 5B and 5D). The cell hyperpolarizes to potentials intermediate between those of E_K_ and E_Cl_ (Fig 5C) driving a net loss of KCl. As the cells dehydrate, the intracellular chloride concentration progressively declines (Fig 8A), driving a net entry of chloride and protons through the Jacobs-Stewart mechanism, with cell acidification and medium alkalinisation (Figs 6A and 8D). Concurrently, dehydration progressively increases the osmotic contribution of haemoglobin along a power function (Fig 8E), elevating its colloidosmotic strength, causing water retention and the generation of a progressively hypertonic effluent with overall increase in system osmolarity (Fig 6C), higher the higher the haematocrit of the cell suspension (Fig 4G). Water retention, in turn, contributes to intracellular chloride dilution, an additional contributor to the overall pH change via rA.

**Figure 9.**
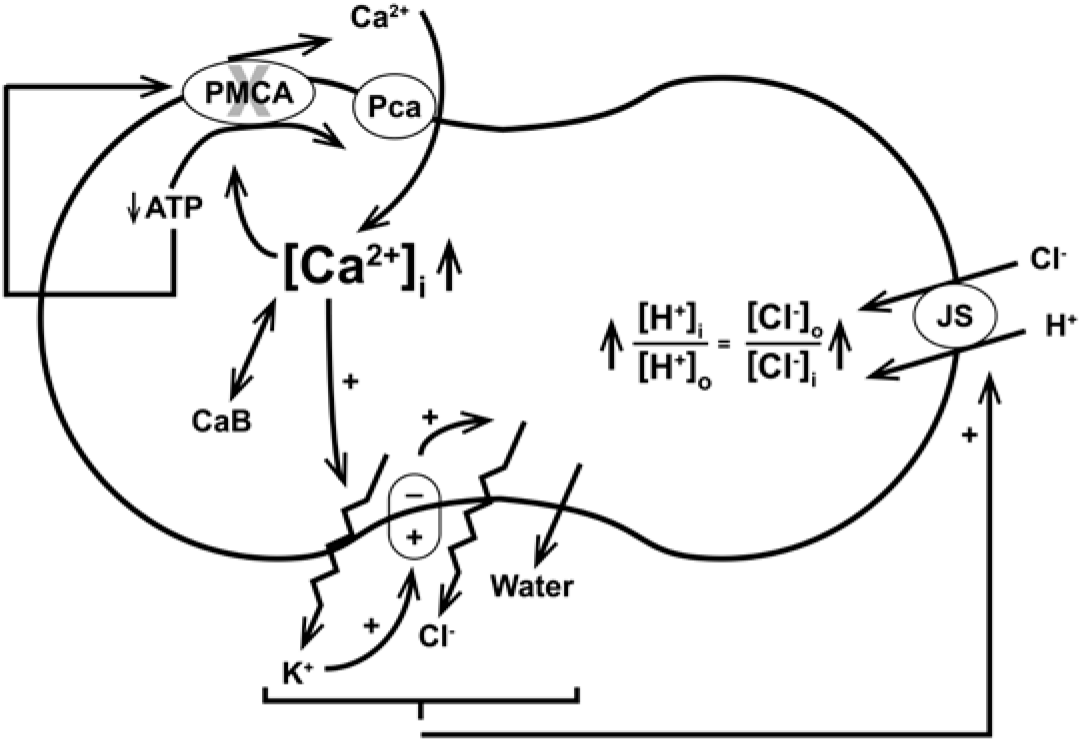
Illustration of the sequence of effects triggered by the Gardos effect, leading from ATP depletion to cell dehydration and acidification. Full description in the text.

## APPENDIX

I. Glossary of variable names as in the column sequence of the model output files (*.csv), with brief description and units
II. Modelling framework flowchart
III. User interface for building simulated experimental protocols
IV. HELP pages: Reference State, Dynamic State and PIEZO1 routine

## I. Glossary of variable names as in the column sequence of the model output files (*.csv), with brief description and units

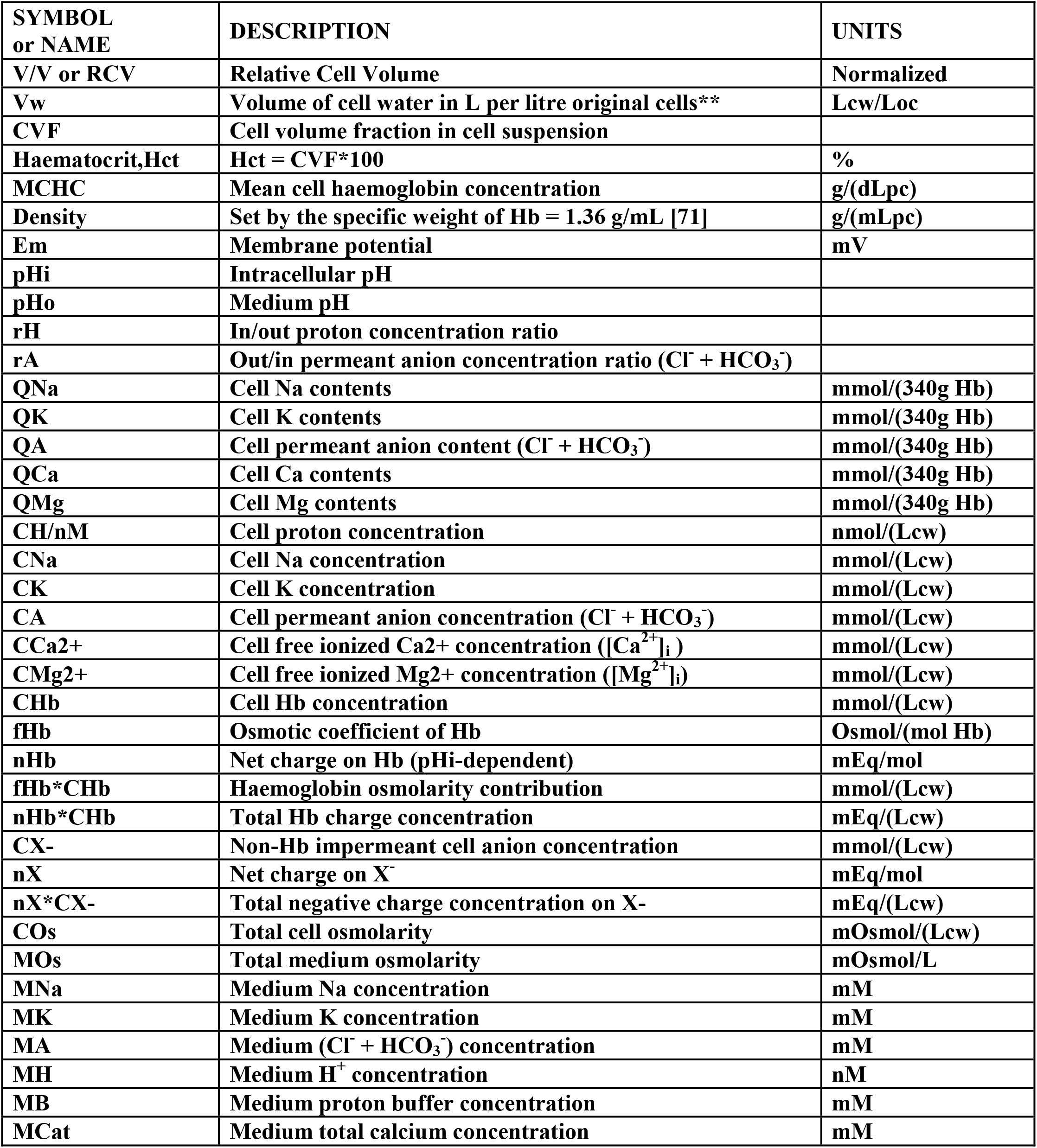
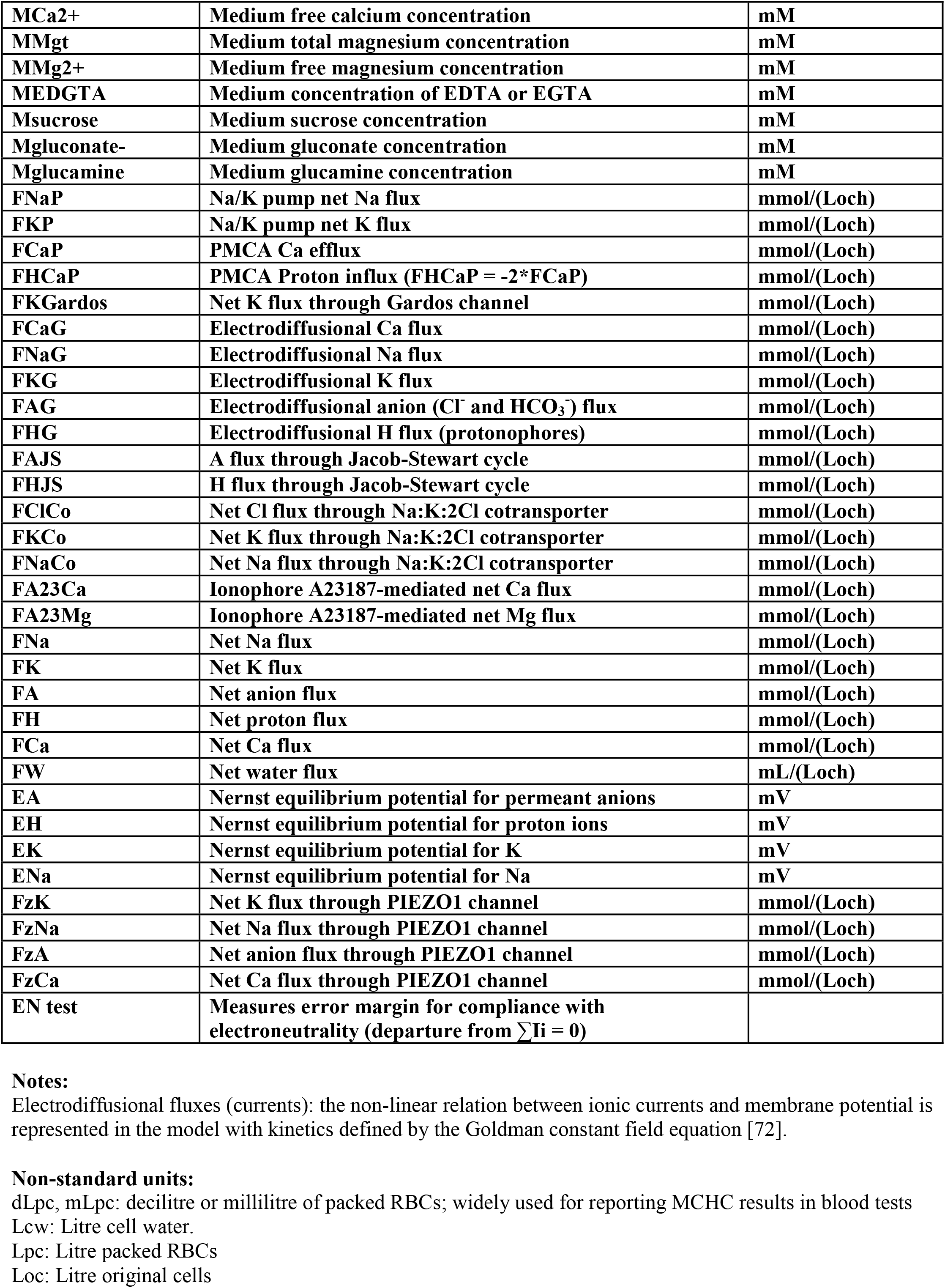
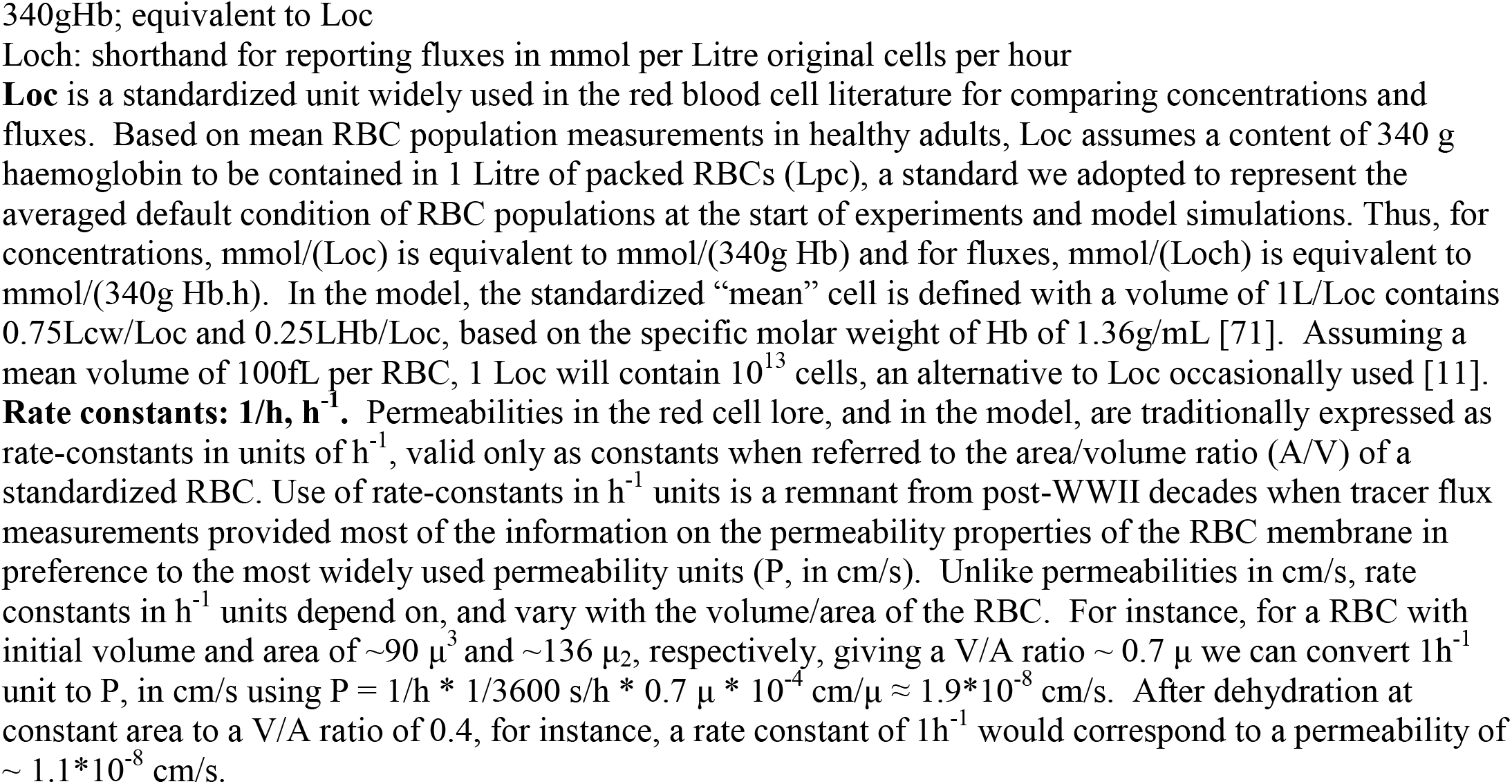

## II. Modelling framework flowchart

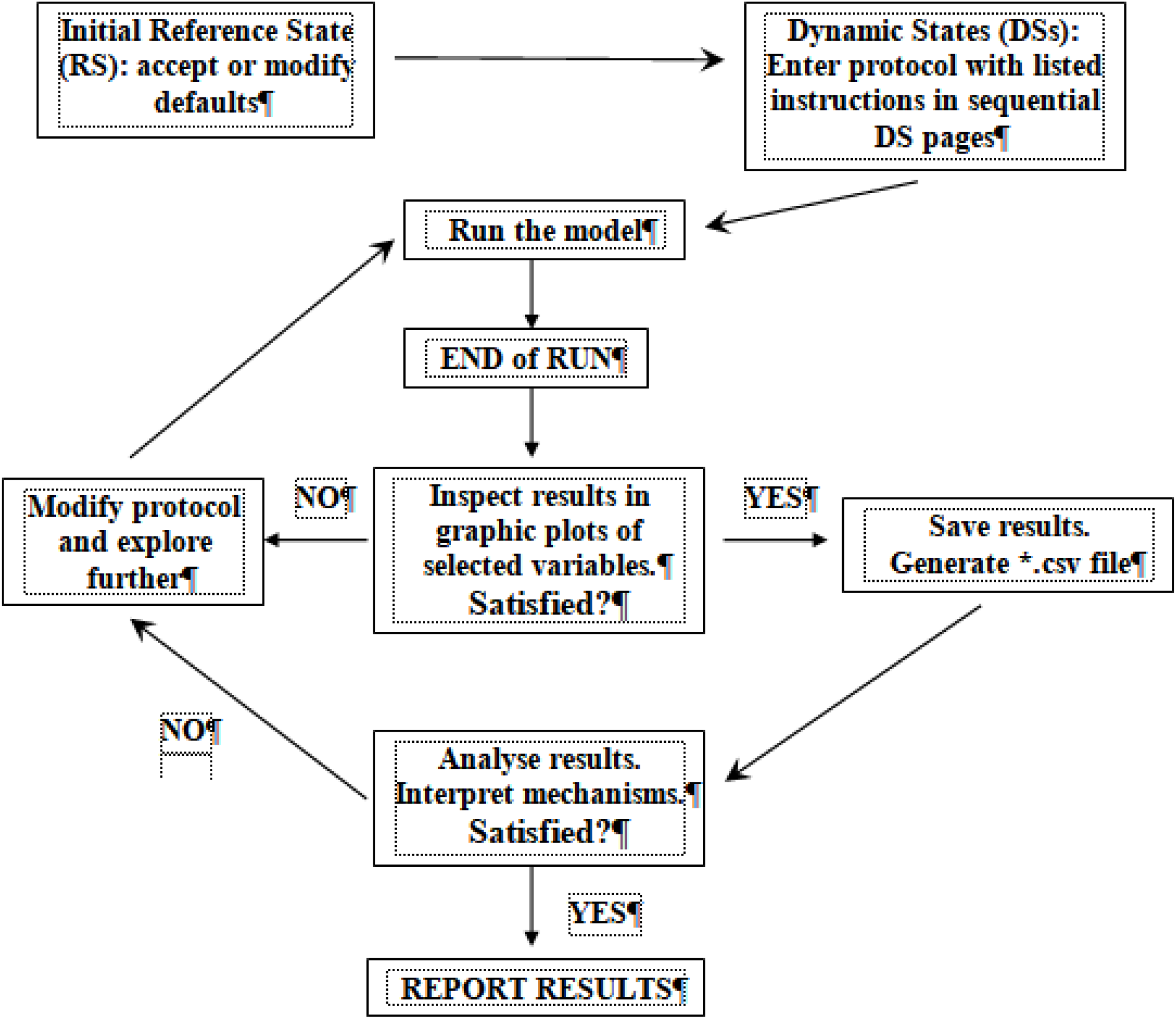

## III. User interface for building simulated experimental protocols

The following Tables list all the entries in Reference (RS) and Dynamic States (DS), with names, brief description of their function, default numerical value, and units. This is how the information appears on the different RS and DS windows. For variables, the same names are used as column headings in the csv output file.

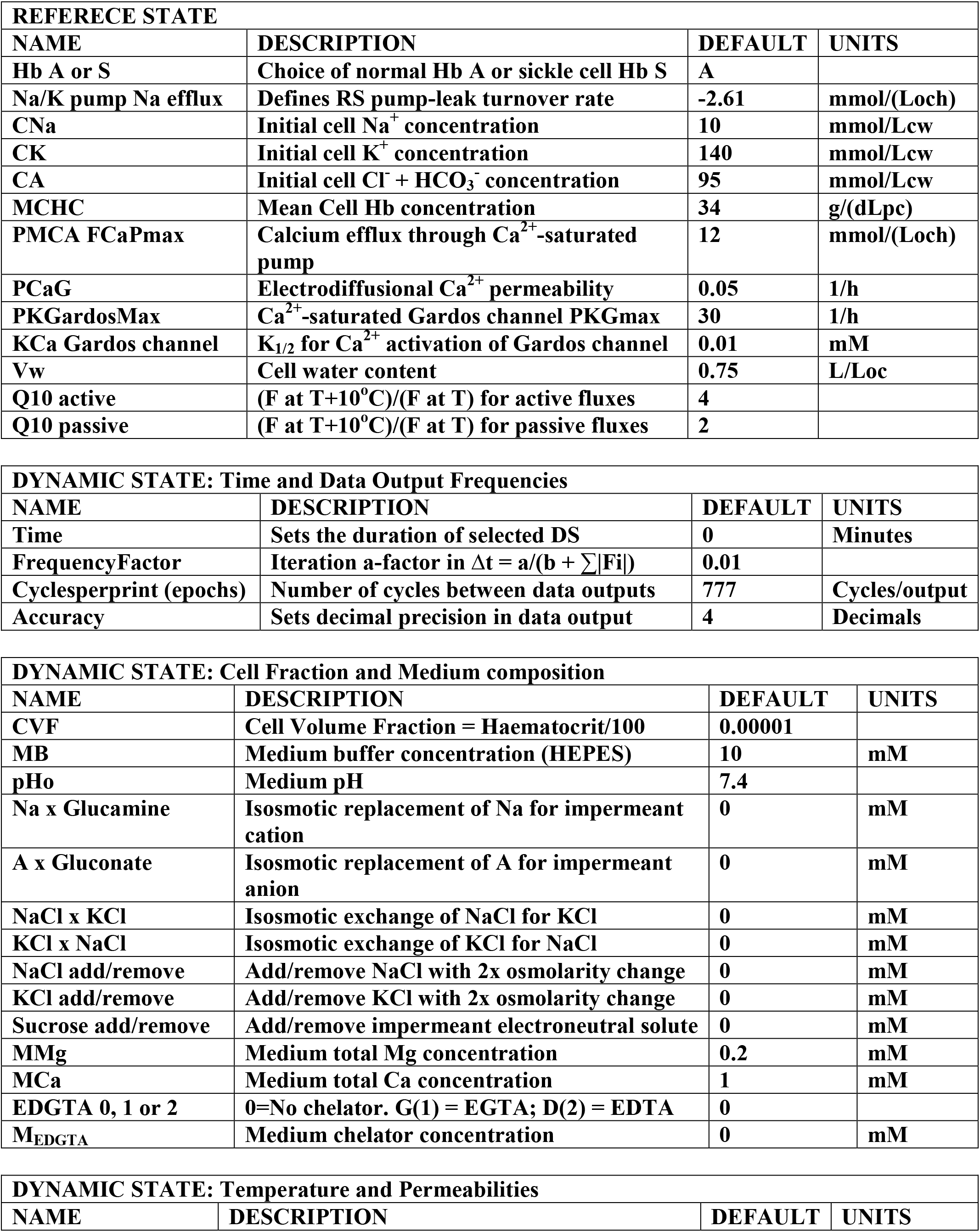
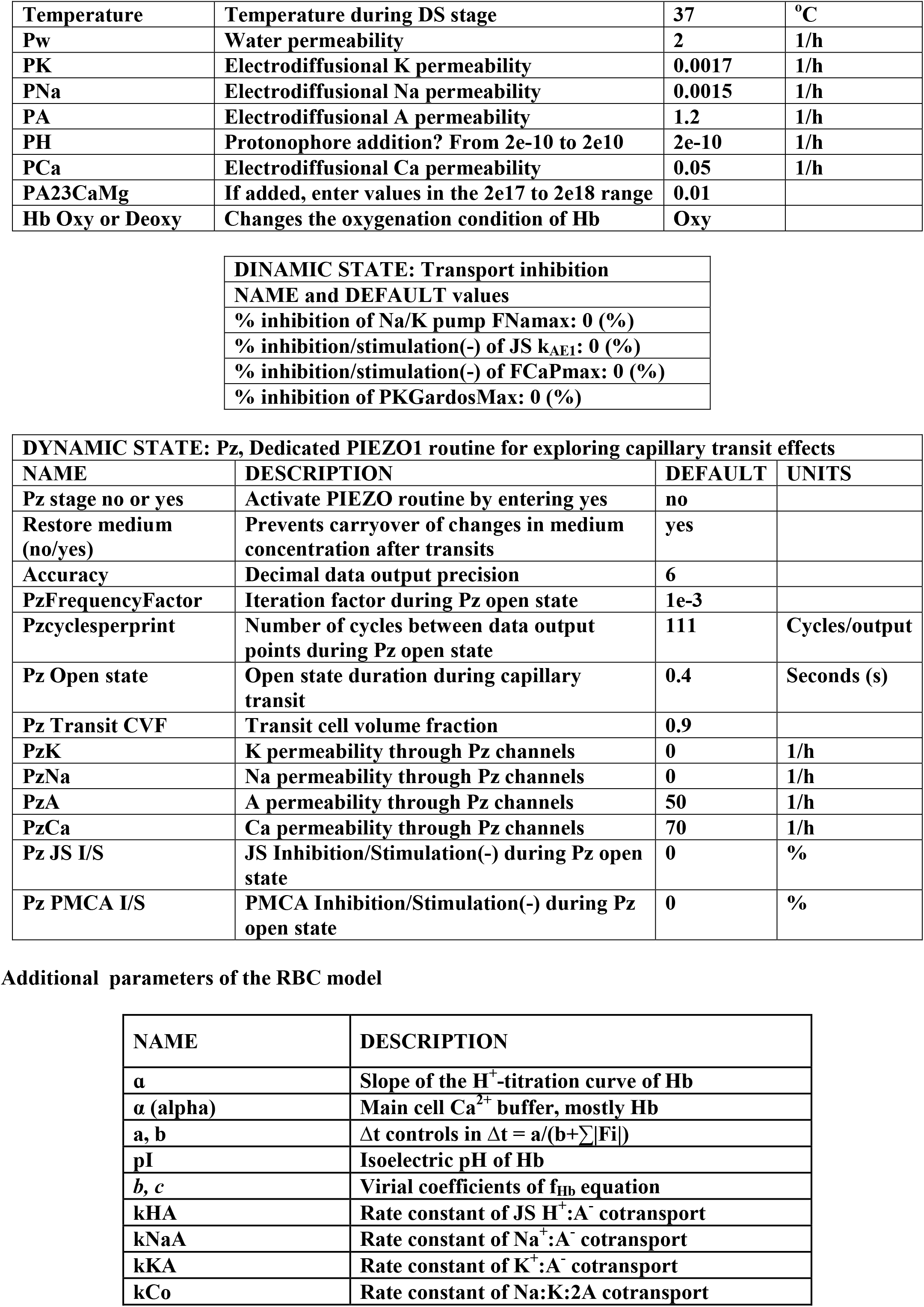
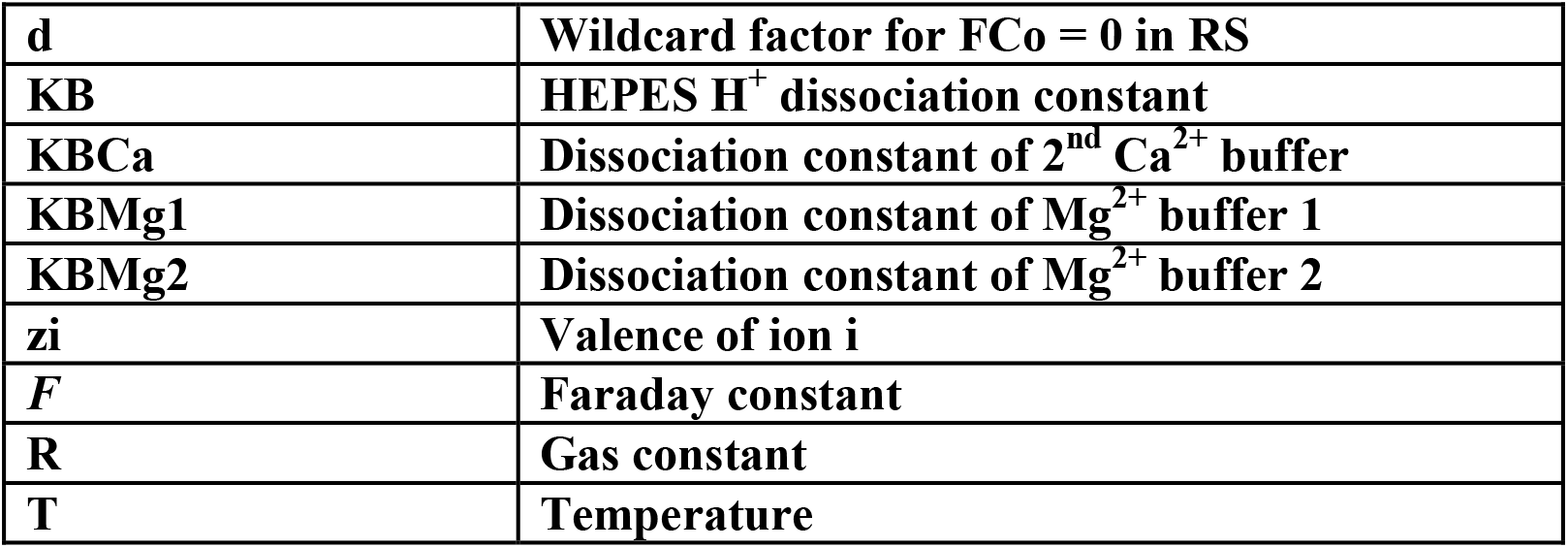

| REFERENCE STATE |  |  |  |
| --- | --- | --- | --- |
| NAME | DESCRIPTION | DEFAULT | UNITS |
| Hb A or S | Choice of normal Hb A or sickle cell Hb S | A |  |
| Na/K pump Na efflux | Defines RS pump-leak turnover rate | -2.61 | mmol/(Loch) |
| CNa | Initial cell Na<sup>+</sup> concentration | 10 | mmol/Lcw |
| CK | Initial cell K<sup>+</sup> concentration | 140 | mmol/Lcw |
| CA | Initial cell Cl<sup>-</sup> + HCO<sub>3</sub><sup>-</sup> concentration | 95 | mmol/Lcw |
| MCHC | Mean Cell Hb concentration | 34 | g/(dLpc) |
| PMCA FCaPmax | Calcium efflux through Ca<sup>2+</sup>-saturated pump | 12 | mmol/(Loch) |
| PCaG | Electrodiffusional Ca<sup>2+</sup> permeability | 0.05 | 1/h |
| PKGardosMax | Ca<sup>2+</sup>-saturated Gardos channel PKGmax | 30 | 1/h |
| KCa Gardos channel | K<sub>1/2</sub> for Ca<sup>2+</sup> activation of Gardos channel | 0.01 | mM |
| Vw | Cell water content | 0.75 | L/Loc |
| Q10 active | (F at T+10°C)/(F at T) for active fluxes | 4 |  |
| Q10 passive | (F at T+10°C)/(F at T) for passive fluxes | 2 |  |

| DYNAMIC STATE: Time and Data Output Frequencies |  |  |  |
| --- | --- | --- | --- |
| NAME | DESCRIPTION | DEFAULT | UNITS |
| Time | Sets the duration of selected DS | 0 | Minutes |
| FrequencyFactor | Iteration a-factor in $\Delta t = a/(b + \sum \vert F_i\vert )$ | 0.01 | |
| Cyclesperprint (epochs) | Number of cycles between data outputs | 777 | Cycles/output |
| Accuracy | Sets decimal precision in data output | 4 | Decimals |

| DYNAMIC STATE: Cell Fraction and Medium composition |  |  |  |
| --- | --- | --- | --- |
| NAME | DESCRIPTION | DEFAULT | UNITS |
| CVF | Cell Volume Fraction = Haematocrit/100 | 0.00001 |  |
| MB | Medium buffer concentration (HEPES) | 10 | mM |
| pHo | Medium pH | 7.4 |  |
| Na x Glucamine | Isosmotic replacement of Na for impermeant cation | 0 | mM |
| A x Gluconate | Isosmotic replacement of A for impermeant anion | 0 | mM |
| NaCl x KCl | Isosmotic exchange of NaCl for KCl | 0 | mM |
| KCl x NaCl | Isosmotic exchange of KCl for NaCl | 0 | mM |
| NaCl add/remove | Add/remove NaCl with 2x osmolarity change | 0 | mM |
| KCl add/remove | Add/remove KCl with 2x osmolarity change | 0 | mM |
| Sucrose add/remove | Add/remove impermeant electroneutral solute | 0 | mM |
| MMg | Medium total Mg concentration | 0.2 | mM |
| MCa | Medium total Ca concentration | 1 | mM |
| EDGTA 0, 1 or 2 | 0=No chelator. G(1) = EGTA; D(2) = EDTA | 0 |  |
| M<sub>EDGTA</sub> | Medium chelator concentration | 0 | mM |

| DYNAMIC STATE: Temperature and Permeabilities |  |  |  |
| --- | --- | --- | --- |
| NAME | DESCRIPTION | DEFAULT | UNITS |
| **Temperature** | **Temperature during DS stage** | **37** | **°C** |
| **Pw** | **Water permeability** | **2** | **1/h** |
| **PK** | **Electrodiffusional K permeability** | **0.0017** | **1/h** |
| **PNa** | **Electrodiffusional Na permeability** | **0.0015** | **1/h** |
| **PA** | **Electrodiffusional A permeability** | **1.2** | **1/h** |
| **PH** | **Protonophore addition? From 2e-10 to 2e10** | **2e-10** | **1/h** |
| **PCa** | **Electrodiffusional Ca permeability** | **0.05** | **1/h** |
| **PA23CaMg** | **If added, enter values in the 2e17 to 2e18 range** | **0.01** |  |
| **Hb Oxy or Deoxy** | **Changes the oxygenation condition of Hb** | **Oxy** |  |

|  |
| --- |
| **DINAMIC STATE: Transport inhibition** |
| **NAME and DEFAULT values** |
| **% inhibition of Na/K pump F<sub>N</sub>amax: 0 (%)** |
| **% inhibition/stimulation(-) of JS k<sub>AE1</sub>: 0 (%)** |
| **% inhibition/stimulation(-) of F<sub>Ca</sub>Pmax: 0 (%)** |
| **% inhibition of PKGardosMax: 0 (%)** |

| **DYNAMIC STATE: Pz, Dedicated PIEZO1 routine for exploring capillary transit effects** |  |  |  |
| --- | --- | --- | --- |
| **NAME** | **DESCRIPTION** | **DEFAULT** | **UNITS** |
| **Pz stage no or yes** | **Activate PIEZO routine by entering yes** | **no** |  |
| **Restore medium (no/yes)** | **Prevents carryover of changes in medium concentration after transits** | **yes** |  |
| **Accuracy** | **Decimal data output precision** | **6** |  |
| **PzFrequencyFactor** | **Iteration factor during Pz open state** | **1e-3** |  |
| **Pzcyclesperprint** | **Number of cycles between data output points during Pz open state** | **111** | **Cycles/output** |
| **Pz Open state** | **Open state duration during capillary transit** | **0.4** | **Seconds (s)** |
| **Pz Transit CVF** | **Transit cell volume fraction** | **0.9** |  |
| **PzK** | **K permeability through Pz channels** | **0** | **1/h** |
| **PzNa** | **Na permeability through Pz channels** | **0** | **1/h** |
| **PzA** | **A permeability through Pz channels** | **50** | **1/h** |
| **PzCa** | **Ca permeability through Pz channels** | **70** | **1/h** |
| **Pz JS I/S** | **JS Inhibition/Stimulation(-) during Pz open state** | **0** | **%** |
| **Pz PMCA I/S** | **PMCA Inhibition/Stimulation(-) during Pz open state** | **0** | **%** |

Additional parameters of the RBC model
| **NAME** | **DESCRIPTION** |
| --- | --- |
| **$\alpha$** | **Slope of the H<sup>+</sup>-titration curve of Hb** |
| **$\alpha$ (alpha)** | **Main cell Ca<sup>2+</sup> buffer, mostly Hb** |
| **a, b** | **$\Delta t$ controls in $\Delta t = a/(b + \sum \vert F_i\vert )$** |
| **pI** | **Isoelectric pH of Hb** |
| **b, c** | **Virial coefficients of f<sub>Hb</sub> equation** |
| **kHA** | **Rate constant of JS H<sup>+</sup>:A<sup>-</sup> cotransport** |
| **kNaA** | **Rate constant of Na<sup>+</sup>:A<sup>-</sup> cotransport** |
| **kKA** | **Rate constant of K<sup>+</sup>:A<sup>-</sup> cotransport** |
| **kCo** | **Rate constant of Na:K:2A cotransport** |
| **d** | **Wildcard factor for FCo = 0 in RS** |
| **KB** | **HEPES H<sup>+</sup> dissociation constant** |
| **KBCa** | **Dissociation constant of 2<sup>nd</sup> Ca<sup>2+</sup> buffer** |
| **KBMg1** | **Dissociation constant of Mg<sup>2+</sup> buffer 1** |
| **KBMg2** | **Dissociation constant of Mg<sup>2+</sup> buffer 2** |
| **zi** | **Valence of ion i** |
| **F** | **Faraday constant** |
| **R** | **Gas constant** |
| **T** | **Temperature** |

## IV. HELP pages: (1) The Reference State, (2) The Dynamic State, and (3) The PIEZO1 subroutine

## (1) The Reference State (RS)

The RS is a steady-state representing the initial condition of a RBC at the start of experiments. Changing default values automatically recalculates a new initial steady-state with equations that ensure compliance with charge and osmotic equilibrium within defined error margins. To approximate the condition of a young RBC, for instance, one could replace the corresponding defaults by CNa 5, CK 145, Vw 0.85 and FNaP - 3.2.

**HbA or HbS:** HbA and HbS differ in the values of their isoelectric point and net charge per mol, *a*. For HbA (default) pI(0°C) = 7.2 and *a* = −10 Eq/(mol*Δ(pH – pI) unit); for HbS, the corresponding values are 7.4 and −8. The net charge on Hb at each pHi, n_Hb_, is computed from n_Hb_ = *a\**(pHi – pI), where *a* is the slope of the proton titration curve of Hb in Eq/mol, and pI is the isoelectric pH of Hb. The model computes the change in pI with temperature (T) following: pI(T°C) = pI(0°C) - 0.016*T(°C) rendering a pI at 37°C for HbA of about 6.6 [11].

**Na/K pump Na efflux (FNaP):** Changing the Na flux automatically resets the associated pump-mediated K influx and reverse Na/K fluxes following the stoichiometries and relative forward-reverse flux ratio encoded in the model.

**CNa and CK:** Initial Na^+^ and K^+^ concentrations in cell water (mmol/Lcw)

**CA:** Initial cell Cl^−^ + HCO_3_^−^ concentration (mmol/Lcw).

**MCHC:** “Mean Cell Haemoglobin Concentration”, a common haematological parameter in blood test assays, traditionally reported in gHb/dLoc; MCHC of “mean” model cell = 34 gHb/dLpc

**PMCA Fmax (FCaPmax):** Maximal Ca^2+^ extrusion flux through an ATP and Ca^2+^-saturated plasma membrane calcium pump

**PKGardosMax:** electrodiffusional K^+^ permeability through Ca^2+^-saturated Gardos channels

**KCa Gardos channel:** Half-maximal Ca^2+^ dissociation constant (K_1/2_) for Gardos channel activation

**Vw:** Water content associated with 340 g Hb; the volume occupied by 340 g Hb tetramer with a molar weight of 1.36 g/ml is 0.25 L. The default 0.75 Lcw/Loc for Vw sets a value of 1 L/Loc for the initial volume of the default “reference” RBC.

**Q10 active or passive:** The Q10 factors determine the extent by which active and passive fluxes (F) are set to increase or decrease for each 10°C increase or decrease in temperature, T

## (2) The Dynamic State (DS)

Displays five tabs grouping lists of parameters and variables with default values for constructing one or successive stages in simulated experimental protocols.

## Time &Data Output Frequency

**Time:** Sets the duration of each DSn stage

**FrequencyFactor:** Sets the duration of each iteration interval (Δt at time = t) inversely proportional to the absolute value of the sum of all the net fluxes across the cell membrane at time t. Allows data output frequencies to appear proportional to the rate of change in the system at constant “cyclesperprint” values.

**Cyclesperprint:** sets the number of computational cycles between data output points

**Accuracy:** sets the decimal precision on the output data

## Cell fraction and Medium Composition

Medium concentrations of X are indicated by MX in mM units. Isosmotic exchanges of X for Y are shown as X x Y. Addition/removals allow changes in medium osmolarity. HEPES, Glucamine, gluconate, sucrose, Mg, EGTA and EDTA are treated as impermeant medium solutes. Sucrose represents any electroneutral impermeant small solute used to alter medium osmolarity. Gluconate and glucamine represent any impermeant monovalent ion used to replace A or Na in the medium.

**MB:** Medium buffer concentration, HEPES by default

**EDGTA 0, 1, 2:** Prompts for the addition of EGTA (G(1)) or EDTA (D(2)) to the cell suspension.

**M_EDGTA_:** Prompts for the concentration of EGTA or EDTA, if added. The default is 0, no addition.

## Temperature & Permeabilities

**Notation on the unit used for electrodiffusional ion permeabilities, 1/h or h^−1^:** 1/h is a widely used permeability unit in the RBC literature. For permeability comparisons between membranes from different cell types the most widely used unit is cm/s. For RBCs, both units are related through P_cm/s_ = P_1/h_*(V/A)/3600, where V and A correspond to the RBC volume and membrane area (in cm units) at the time the permeability measurement was taken.

**PHG:** PHG was modelled to enable simulations of the effects of protonophore additions. The default value represents no protonophore present. To simulate observed effects change PHG from 2e-1- to 2e10.

**PA23CaMg:** Ionophore A23187 mediates electroneutral X^2+^:2H^+^ exchanges with well defined highly non-linear kinetics in human RBCs. The default value represents absence of ionophore. To simulate the effects of ionophore concentrations capable of generating a Ca^2+^ flux exceeding that of the PMCA Fmax at medium Ca^2+^ concentrations around 0.2 mM use values around 2e18.

**Hb pI(0°C) oxy(7.2), deoxy(7.5):** Hb is assumed to be in a oxygen-saturated condition by default (Oxy).

**Deoxygenation of Hb** (Deoxy) changes its pI(0°C) from 7.2 to 7.5. The model automatically adjusts the actual pI change for the temperature of the experiment. The pI shifts during oxy-deoxy transitions cause changes in the protonization condition of Hb with secondary changes in pHi and [Mg^2+^]_i_, changes which the model predicts with verified accuracy.

## Transport inhibition/stimulation (%; defaults = 0)

The default Fmax value for each transporter, Fm, is modified according to Fm*(100-X)/100 where X is the % inhibition entered at the prompt. Fm stays modified in successive DS stages unless modified again. Entries in successive DSs always apply to the original default (0%). To return to the original uninhibited Fm enter “0”. The same equation delivers stimulation if you enter negative numbers. It applies to JS and PMCA entries only. For an n-fold stimulation, enter “−n*100”, for instance “−200” for a two-fold stimulation. In the model, the intrinsic electrodiffusional anion permeability of the RBC, PA, is treated as a permeability pathway independent of the anion exchanger, thus preventing prejudged linkage associations when implementing the JS inhibition/stimulation subroutine [18].

## (3) The PIEZO1 routine

All the membrane transporters represented in the red cell model are active all the time and participate to different extents in the dynamic responses to perturbations. PIEZO1 is unique in that it is generally inactive by default, responding only to conditions eliciting cell deformation, for instance during capillary transits. The dedicated PIEZO tag allows exploration of the potential effects of PIEZO1 activation during capillary transits. Preliminary tests showed that because of the brief duration of each transit (less than 1s), the magnitude of the changes in model variables was extremely small even attributing extremely high permeabilities to PIEZO1 mediation. This required a substantial increase in decimal accuracy and in the density of data acquisition. The tag offers a protocol design with defaults open to change by the user, as outlined next.

I. In a DS selected for PIEZO1 implementation start by assigning to it a duration of 5min (Time: 5 (min))
II. Double click on the “Incorporate PIEZO stage: no” and enter “yes”
III. This brings up a predesigned DS stage with the following defaults or assignments:

1. Incorporate PIEZO stage: yes
2. Pz Restore Medium: Yes
3. Accuracy: 6
4. Pz frequency factor: 1e-3
5. Pz cycles per print: 111
6. Pz Open state: 0.4 (s)
7. Pz Transit cell volume fraction: 0.9
8. PzK: 0 (1/h)
9. PzNa: 0 (1/h)
10. PzA: 50 (1/h)
11. PzCa: 70 (1/h)
12. Pz JS I/S: 0 (%)
13. Pz PMCA I/S: 0 (%)

Pz identifies PIEZO-related parameters and variables. The default five-minute duration of the PIEZO1 DS stage covers three consecutive periods: an initial two-minute baseline control, a PIEZO1 open state period, and a PIEZO1 closure period for the remainder time to five minutes to allow assessment of the short-term reversibility of the changes induced during the open state. Pz accuracy determines the decimal precision of data output in the csv file. Pz frequency factor and Pz cycles per print determine the density of data output over this period. The PzX values list the electrodiffusional permeabilities assigned by default to K^+^, Na^+^, A^−^, and Ca^2+^ in the open state. The precise ion selectivity, conductance and inactivation kinetics with which PIEZO1 channels operate in human RBCs is unknown at present. The default permeability values chosen render model outcomes in conformity with the scarce but solid experimental evidence available in RBCs [57, 58, 73], as extensively analysed and discussed in the first application of the PIEZO1 model extension to investigate the circulatory physiopathology of human RBCs [74, 75]. The value of 0.9 used for the cell volume fraction applies strictly for the duration assigned to the capillary transit, to approximate the squeezed transit condition of the RBCs between the highly diluted (CFV ~ 0.00001) conditions in the systemic circulation before and after transits. The infinite reservoir composition of the medium representing the systemic circulation is preserved by the Restore Medium subroutine. The Pz JS and PMCA Pz Inhibition/Stimulation options enable study of the effects of anion-proton equilibration rates through the Jacobs-Stewart cycle, and of different levels of PMCA activity, but only during Pz open states. The options within the Transport inhibition tag operate only for periods outside Pz open states.

